# BubR1 recruitment to the kinetochore via Bub1 enhances Spindle Assembly Checkpoint signaling

**DOI:** 10.1101/2021.08.12.456107

**Authors:** Anand Banerjee, Chu Chen, Lauren Humphrey, John J. Tyson, Ajit P. Joglekar

## Abstract

During mitosis, unattached kinetochores in a dividing cell activate the Spindle Assembly Checkpoint (SAC) and delay anaphase onset by generating the anaphase-inhibitory Mitotic Checkpoint Complex (MCC). These kinetochores generate the MCC by recruiting MCC constituent proteins, including BubR1. In principle, BubR1 recruitment to signaling kinetochores should increase its local concentration and promote MCC formation. However, in human cells BubR1 is mainly thought to sensitize the SAC to silencing; whether BubR1 localization to signaling kinetochores per se enhances SAC signaling activity remains unknown. Therefore, we used ectopic SAC activation systems (eSAC) to isolate two molecules that recruit BubR1 to the kinetochore: the checkpoint protein Bub1 and the KI and MELT motifs in the kinetochore protein KNL1, and observed their contribution to eSAC signaling. Our quantitative analyses and mathematical modeling show that the Bub1-mediated BubR1 recruitment to the human kinetochore promotes SAC signaling and highlight BubR1’s dual role of directly strengthening the SAC and indirectly silencing it.

## Introduction

The “Spindle Assembly Checkpoint” (SAC) is a cell cycle control that minimizes chromosome missegregation during cell division (Lara-Gonzalez et al., 2021b; Musacchio, 2015). It is activated by kinetochores that are not stably attached to the plus-ends of spindle microtubules. These kinetochores generate a diffusible anaphase-inhibitory signal known as the Mitotic Checkpoint Complex (MCC). The MCC delays anaphase onset to avert cell division in the presence of unattached kinetochores. The rate at which an unattached kinetochore generates MCC depends on its ability to recruit SAC signaling proteins, which include constituent proteins of the MCC: Bub1-Bub3, BubR1-Bub3, Mad1-Mad2, and Cdc20 (Figure 1A, dashed square). This signaling cascade generates either the MCC itself, its sub-complex C-Mad2-Cdc20, or both. Given this knowledge, it is reasonable to expect that the rate of MCC generation at a kinetochore will increase with higher recruitment of signaling proteins, and especially the MCC components, to the kinetochore (Collin et al., 2013; Lara-Gonzalez et al., 2021a; Piano et al., 2021). Interestingly, however, this expectation appears to not hold true for BubR1. Disruption of BubR1 recruitment to unattached kinetochores does not reduce the duration of SAC-induced mitotic arrest in nocodazole-treated cells (Overlack et al., 2015; Zhang et al., 2015). This is because BubR1 recruits to the kinetochore Protein Phosphatase 2A (PP2A), which promotes SAC silencing (Foley et al., 2011; Nijenhuis et al., 2014; Qian et al., 2017). Despite this knowledge, it is crucial to determine whether BubR1 recruitment to the kinetochore promotes SAC signaling by enhancing MCC assembly. This BubR1 activity can be crucial for minimizing chromosome missegregation during normal cell division wherein a small number of unattached kinetochores must activate the SAC and delay anaphase onset (Roy et al., 2020).

**Figure 1.**
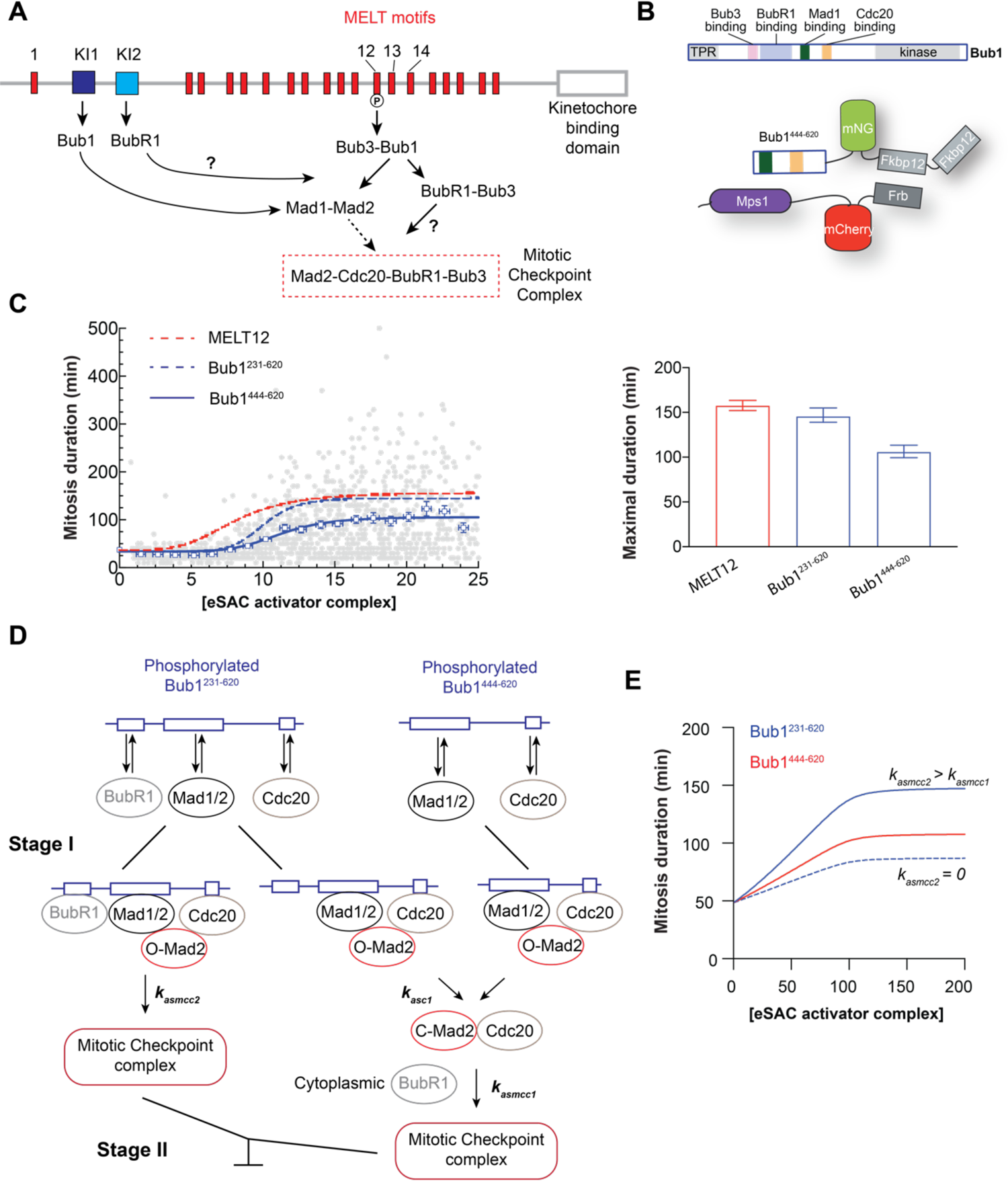
The contribution of Bub1-BubR1 heterodimerization to MCC assembly in eSAC signaling. **A** Schematic at the top displays the domain organization of human Knl1. Below is a simplified schematic displaying the pathways that recruit SAC signaling proteins to unattached kinetochores. **B** Schematic at the top displays the domain organization of human Bub1. The cartoon represents the ectopic SAC activation (eSAC) system. **C** Dose-response data for an eSAC system that uses Bub1^444-620^ as the phosphodomains. Each gray circle represents a single cell (n = 1920 from 2 experiments; 139 and 155 are the lower and upper 95% confidence intervals). The blue squares represent the mean values of the dose-response data binned according to mCherry fluorescence. Vertical and horizontal lines represent SEM. The solid blue curve displays the least squares 4-parameter sigmoidal fit to the binned mean values. Hill equation fits for dose-response data for one MELT motif (MELT12, dashed red curve) and an extended Bub1^231-620^ phosphodomain containing the Bub3- and BubR1-binding sites (dashed blue curve) from our previous study have been super-imposed for comparison (Chen et al., 2019). The bar graph on the right displays the maximal mitotic duration estimated by the fit. Error bars represent 95% confidence intervals of the fitted maximal mitotic duration. **D** Cartoon displays the simplified scheme used to simulate the generation of MCC by the eSAC system. **E** Numerical simulation of the dose-response curves by assuming the assembly of signaling complexes and ultimately the MCC as shown in **D** followed by a model described by He et al (He et al., 2011).

In this study, we examine the contribution of BubR1 recruitment to kinetochore-mediated SAC signaling. We first use the ectopic SAC activation system or the ‘eSAC’ to find that the binding of BubR1 to Bub1 elevates Bub1-mediated MCC generation (Chen et al., 2019). On the other hand, the recruitment of Bub1 and BubR1 via the ‘KI’ motifs in the kinetochore protein Knl1 does not contribute to MCC generation mediated by the MELT motifs within the Knl1 phosphodomain (Bolanos-Garcia et al., 2011a; Bolanos-Garcia et al., 2011b; Kiyomitsu et al., 2011; Krenn et al., 2012; Primorac et al., 2013; Vleugel et al., 2013; Zhang et al., 2014). We also establish a mathematical model to elucidate the mechanistic details of the SAC signaling cascade that generates the MCC. Finally, we also demonstrate that BubR1 recruitment to the kinetochore via Bub1 promotes SAC signaling.

## Results & Discussion

### The BubR1 binding domain of Bub1 promotes Bub1-mediated MCC assembly

Bub1 coordinates the rate-limiting step in MCC assembly: the formation of the closed-Mad2-Cdc20 (Mad2:Cdc20) sub-complex (Faesen et al., 2017; Lara-Gonzalez et al., 2021a; Piano et al., 2021). Mad2:Cdc20 must bind BubR1 to complete MCC formation. BubR1 is recruited to the kinetochore by Bub1, and the KI motifs and MELT motifs in KNL1 (Bolanos-Garcia et al., 2011b; Kiyomitsu et al., 2011; Overlack et al., 2015; Zhang et al., 2016). Whether this BubR1 recruitment promotes MCC formation remains unclear. In fission yeast, BubR1 binding to Bub1 is essential for SAC activity (Leontiou et al., 2019). However, BubR1 is unlikely to be recruited to budding yeast kinetochores (Roy et al., 2022; Tromer et al., 2016), and in nocodazole-treated HeLa cells, the disruption of Bub1-mediated BubR1 recruitment slightly strengthens the SAC (Overlack et al., 2015; Zhang et al., 2016) because the disruption of BubR1 recruitment also disrupted BubR1-mediated PP2A recruitment (Elowe et al., 2007; Suijkerbuijk et al., 2012). Therefore, to isolate and quantitatively define the effect of Bub1-BubR1 heterodimerization on Bub1-mediated MCC assembly, we used the ectopic SAC activation system or the eSAC.

Forced dimerization of a fragment of the central domain of Bub1 with Mps1 delays anaphase onset in HeLa cells, budding yeast, and fission yeast (Aravamudhan et al., 2015; Chen et al., 2019; Leontiou et al., 2019; Yuan et al., 2017). In HeLa cells, induced dimerization of the central domain of Bub1 (Bub1^231-620^-mNG-2xFkbp12, diagram in Figure 1B) with the Mps1 kinase domain (Frb-mCherry-Mps1^500-817^) produces a dosage-dependent increase in the duration of mitosis with a maximal duration of ∼ 145 minutes (Figure 1C replotted for comparison from (Chen et al., 2019)). Importantly, the extended mitosis was due to increased Mad2-Cdc20 and MCC formation in HeLa cells and fission yeast (Leontiou et al., 2019; Roy et al., 2022). To assess the contribution of Bub1-BubR1 heterodimerization to Bub1-mediated MCC assembly, we created a truncated Bub1 phosphodomain lacking the BubR1 heterodimerization domain (Bub1^444-620^-mNG-2xFkbp12) (Overlack et al., 2015; Zhang et al., 2016). Rapamycin-induced dimerization of this phosphodomain with Frb-mCherry-Mps1^500-^817 elicited a significantly weaker eSAC activity, with a maximal mitotic duration of 105 ± 6 minutes (Figure 1C, estimated from a fit with the 4-parameter Hill equation, the range indicates 95% confidence intervals, see Methods for details). The absence of the Bub3-binding GLEBS domain in Bub1^444-620^-mNG-2xFkbp12 cannot explain the observed decrease in eSAC activity (Roy et al., 2022). Thus, Bub1-BubR1 heterodimerization promotes MCC mediated by the Bub1 phosphodomain.

### Simulation of the signaling activity of the Bub1 phosphodomains

To quantitatively understand how these events shape the dose-response dependence, we constructed a mathematical model by considering the events at the Bub1 eSAC prior to MCC assembly. This model consists of two stages. In the first stage, we calculate the steady-state concentrations of signaling complexes assembled by the Bub1 eSAC phosphodomains assuming mass action kinetics (Figure 1D, equations 1-4, 5a-j). Bub1^231-620^ recruits BubR1 and Cdc20 independently of its phosphorylation state or the presence of other bound proteins; Bub1^444-620^ recruits only Cdc20 (Di Fiore et al., 2015; Overlack et al., 2015). Both phosphodomains are activated by Mps1-mediated phosphorylation, after which they recruit Mad1-Mad2 (abbreviated as Mad1/2) (Faesen et al., 2017; Ji et al., 2017). Therefore, the signaling activity of each phosphodomain will be proportional to the amount of Frb-mCherry-Mps1^500-817^, i.e., the eSAC dosage, and it will be limited by the cellular Mad1/2 abundance when the eSAC dosage exceeds Mad1/2 abundance (shown later, see Figure S1A). Using these interactions, Bub1^231-620^ can assemble two types of signaling complexes: one that contains Cdc20, BubR1, and Mad1/2, and one containing only Cdc20 and Mad1/2; Bub1^444-620^ forms only the signaling complex containing Mad1/2 and Cdc20 (Figure 1D, left).

We simulated MCC formation by the Bub1:BubR1:Mad1/2:Cdc20 signaling complex as follows. Because Mad2-Cdc20 formation is the rate-limiting step in MCC assembly (Faesen et al., 2017), the Bub1^231-620^:BubR1:Mad1/2:Cdc20 signaling complex first assembles Mad2:Cdc20 with the rate constant *k_asc1_*. The newly formed Mad2:Cdc20 can bind BubR1 either within the signaling complex with the rate constant *k_asmcc2_* or in the cytosol with the rate constant *k_asmcc1_* (Figure 1D, bottom). The model assumes that all the Mad2:Cdc20 formed by Bub1^231-620^:BubR1:Mad1/2:Cdc20 binds BubR1 within the signaling complex. However, allowing a reasonable fraction of Mad2:Cdc20 to escape from the signaling complex does not affect the overall behavior of the model. Because Mad2:Cdc20 formation is the rate-limiting step, we reduce the number of free parameters by assuming that *k_asc1_* and *k_asmcc2_* are equal. The cumulative MCC formed by Bub1^231-620^ is the sum of MCC formed from these two processes. All Mad2-Cdc20 generated by Bub1^441-620^:Mad1/2:Cdc20 forms MCC in the cytosol. In the second stage of the model (Figure 1D, equations 5k-n, Methods), the MCC formed by both processes modulates Cyclin B degradation and thereby controls the timing of metaphase-to-anaphase transition (Figure S1B) (Chen et al., 2019; He et al., 2011).

We first simulated the dose-response curve for the Bub1^444-620^ eSAC involving only cytoplasmic MCC assembly. We retained protein concentrations used in the original He et al. model and used reasonable rate constants for Mad2:Cdc20 formation (*k_asc1_*) and cytosolic MCC formation (*k_asmcc1_*, Figure 1E, red curve; see Table S1 for protein concentrations and rate constants used). Next, we considered MCC formation by the Bub1^231-620^ eSAC, which can assemble the MCC within the signaling complex or the cytosol. If BubR1 recruited by the signaling complex does not participate in MCC formation (i.e., *k_asmcc2_* = 0), the simulation produced a dose-response curve with a lower maximal response (dotted blue line in Figure 1E). This is because BubR1 bound to Bub1 cannot participate in any form of MCC assembly. Therefore, the only effect of Bub1-BubR1 heterodimerization is reduced cytosolic BubR1 concentration, and consequently, reduced rate of MCC generation. This prediction is inconsistent with the data. Therefore, the model predicts that BubR1 recruited by Bub1 must promote MCC formation within the eSAC signaling complex. Following this insight, we assumed that *k_asmcc2_* is 10-fold higher than *k_asmcc1_* for cytosolic MCC assembly. With this change, the simulation produced a longer delay in anaphase onset, matching our observations.

This simulation provides two insights. First, it supports the observation that Bub1-BubR1 heterodimerization promotes MCC formation by the eSAC based on the Bub1 phosphodomain. It also indicates that if Bub1-BubR1 heterodimerization does not promote MCC formation, it will lead to BubR1 sequestration and a reduced rate of cytosolic MCC formation. This effect plays a critical role in the experiments that follow.

### KI motifs suppress the signaling strength of the eSAC phosphodomain

The KI motifs, so named because they contain lysine and iso-leucine residues critical for their activities, recruit Bub1 and BubR1 to the human kinetochore (Figure 1A). The first KI motif (KI1) is thought to exclusively bind Bub1, whereas the second motif (KI2) binds BubR1 (Bolanos-Garcia et al., 2011a; Kiyomitsu et al., 2011; Krenn et al., 2012). A prior study concluded that the KI motifs cooperate with the first MELT motif in KNL1 to strengthen SAC signaling (Krenn et al., 2014; Vleugel et al., 2013). However, it is unclear whether the activity of regulatory enzymes (e.g., PLK1, PP2A) bound to Bub1 and BubR1 plays a role in these observations (Jia et al., 2016; Nijenhuis et al., 2014; von Schubert et al., 2015). Therefore, to test whether Bub1 and BubR1 bound to the KI motifs directly promote MCC assembly and delineate the roles of the KI motifs, we developed an eSAC phosphodomain comprising the two KI motifs and a MELT motif (Knl1^160-256^) (Figure 2A). We also created a variant phosphodomain with inactive KI motifs (Krenn et al., 2012). If Bub1 and BubR1 recruitment via the KI motifs enhances MELT motif activity, this will be apparent as increased signaling strength of this new eSAC phosphodomain compared to the variant phosphodomain with inactive KI motifs.

**Figure 2.**
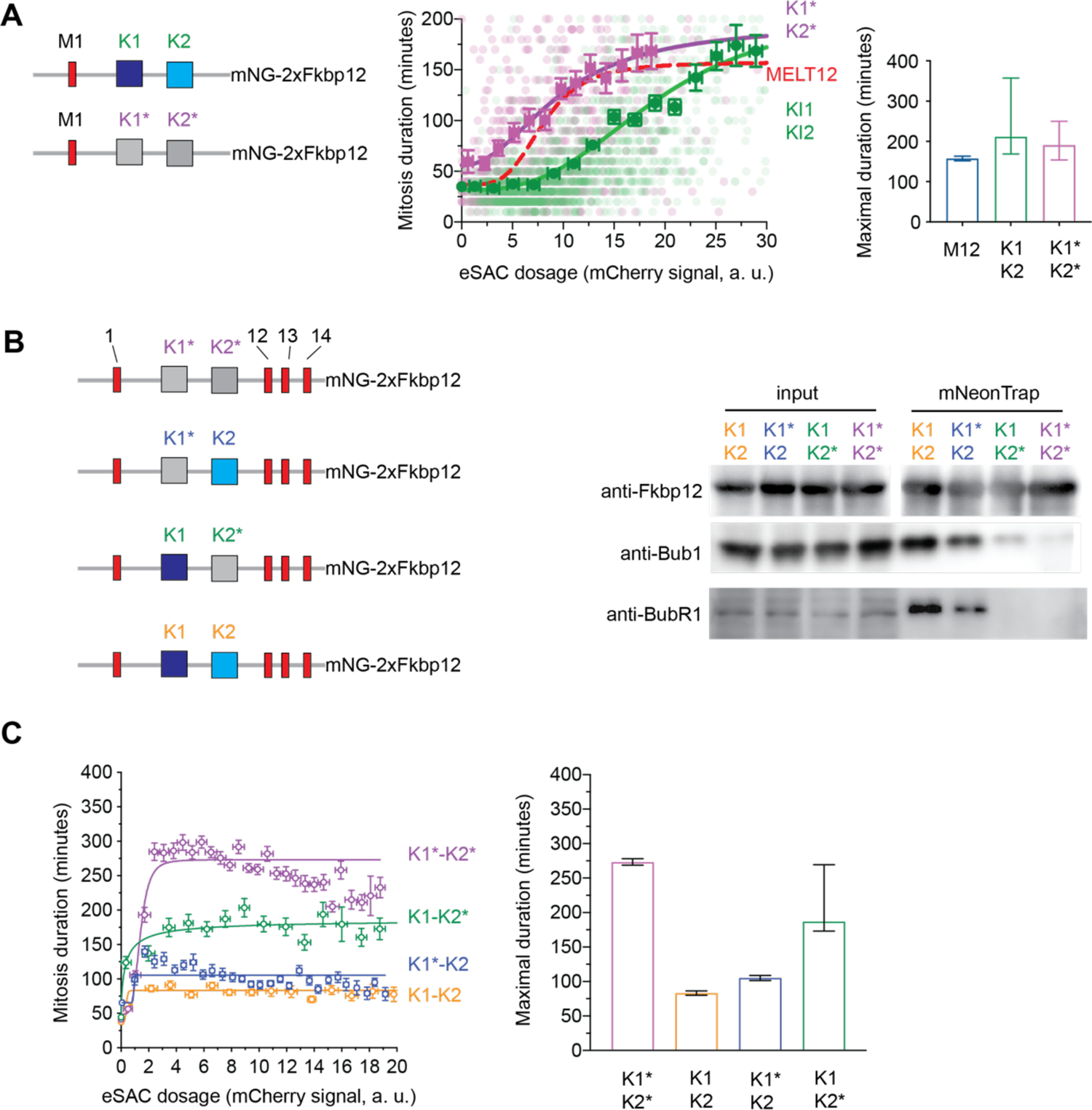
Characterization of the binding of the KI motifs in the eSAC phosphodomain with Bub1 and BubR1 and its contribution to eSAC signaling. **A** Left: Schematic of the two phosphodomains used to test whether the KI motifs contribute to MCC assembly mediated by the MELT motif in the eSAC phosphodomain. KI1-KI2 indicates phosphodomain with intact KI motifs, KI1*-KI2* indicates phosphodomains wherein the KI motifs are inactivated using suitable point mutations (see Methods for details). The scatter plot in the middle displays the dose-response data for the two phosphodomains (n = 1888 for KI1-KI2 (green) and n = 836 for KI1*-KI2* (magenta) from ≥ 2 experiments; symbol usage follows the scheme established in 1**C**). The bar graph on the right displays the maximal mitotic duration predicted by 4-parameter sigmoidal fits to the binned mean values as in Figure 1C. Vertical lines display 95% confidence intervals on the fit parameter. **B** Left: Schematic of the phosphodomains consisting of 4 MELT motifs and either active or inactive (indicated by *) KI motifs. Right: Immunoprecipitation of the eSAC phosphodomains using mNeonGreen-Trap beads followed by immunoblot analysis to probe for the co-IP of the indicated proteins. **C** Dose-response data for the indicated phosphodomains. Only the mean values of binned data are shown for clarity. Data analysis performed as in 1D (n = 1019, 1024, 666, and 3219 from ≥ 2 experiments for KI1-KI2, KI1-KI2*, KI1*-K2, and KI1*-KI2* respectively). The bar graph on the right displays the maximal time in mitosis of the fitted maximal mitotic duration (error bars represent 95% confidence intervals).

We first determined the dependence of mitotic duration on the dosage of the variant phosphodomain with inactive KI motifs (M1-KI1*-KI2*-mNG-2xFkbp12, Figure 2A left). The maximal response for this phosphodomain was higher than the previously defined maximal response for the eSAC phosphodomain containing the 12^th^ MELT motif alone (211 versus 157 minutes with ± 95% confidence intervals of 169-357 and 152-163 respectively predicted by a 4-parameter sigmoidal fit to the binned data). This difference likely results from different Bub1-Bub3 binding affinities of the first and twelfth MELT motif (Chen et al., 2019). Interestingly, active KI motifs in the phosphodomain (M1-KI1-KI2) suppressed its signaling strength: the eSAC concentration required for half-maximal response nearly doubled (EC50 = 9.7 and 19.4 a. u. respectively, Figure 2A middle). The decreased apparent signaling strength of the phosphodomain was eventually compensated by high eSAC dosage as evidenced by the maximal mitotic duration at high eSAC dosage (Figure 2A middle and right). These data suggests that the Bub1 and BubR1 molecules recruited by the KI motifs do not enhance eSAC signaling mediated by the upstream MELT motif.

### Bub1 and BubR1 interactions with the KI motifs do not require Mps1-mediated phosphorylation of KNL1

To better understand the observed suppression of eSAC activity by the KI motifs, we constructed a new phosphodomain by fusing an unstructured region of the Knl1 phosphodomain spanning three previously-characterized MELT motifs (Knl1^881-1014^) to the C-terminus of the phosphodomain used above (Knl1^160-256^ also referred to as ‘M3’, see Figure 2B) (Chen et al., 2019; Vleugel et al., 2015). By incorporating multiple MELT motifs in the phosphodomain, we wanted to test whether the KI motifs influence the ability of the four MELT motifs in the phosphodomain to engage in synergistic signaling (Chen et al., 2019). Based on a previous study, we created mutant phosphodomains wherein the KI motifs were inactive individually or together (KI1*-KI2, KI1-KI2*, and KI1*-KI2*; the asterisk denotes a loss of function, Figure 2B) (Krenn et al., 2012).

We first determined whether the two KI motifs interact with Bub1 and BubR1 exclusively and whether the phosphorylation of MELT motifs is necessary for these interactions. We immunoprecipitated the four mNeonGreen-tagged eSAC phosphodomains from whole-cell extracts of mitotic HeLa cells in the absence of the Mps1 kinase domain using mNeonTrap beads and probed the precipitates for Bub1 and BubR1 (Methods). When both the KI motifs were active (KI1-KI2), Bub1 and BubR1 co-precipitated with the eSAC phosphodomain. As expected, Bub1 and BubR1 did not co-precipitate with the eSAC phosphodomain containing inactive KI motifs (KI1*-KI2*, Figure 2B). Surprisingly, with the first KI motif inactive (KI1*-KI2), which interacts with Bub1 alone, Bub1 still co-precipitated with the phosphodomain, albeit at a lower level. Moreover, BubR1 coprecipitation was reduced. Inactivation of the second KI motif (KI1-KI2*) made BubR1 undetectable in the precipitate and reduced the amount of Bub1. These data are can be explained by the heterodimerization between Bub1 and BubR1, although it remains possible that the second KI motif can interact with Bub1. We obtained similar results from immunoprecipitation experiments involving eSAC phosphodomains with only one MELT motif and the two KI motifs (Supplementary Figure S2A). These experiments show that KI motifs interact with Bub1 and BubR1 constitutively; this interaction does not require the phosphorylation of MELT motifs. They also confirm that the loss-of-function mutants indeed abolish the interactions with Bub1 and BubR1.

An N-terminal fragment of KNL1 spanning residues 1-334 transiently localized to kinetochores, likely by interacting with kinetochore-bound Bub1 or BubR1 or with KNL1 itself (Chen et al., 2019; Kern et al., 2015). Therefore, we examined the localization of the KI1-KI2 and the KI1*-KI2* phosphodomain in mitotic cells treated with Rapamycin using immunofluorescence (Supplementary Figure S2B). Both phosphodomains colocalized with kinetochores, although the amount of kinetochore-localized KI1*-KI2* was ∼ 20% lower than the KI1-KI2 phosphodomain (Supplementary Figure S2B). Despite this localization, the duration of mitosis for the two cell lines in the absence of Rapamycin was similar to the mitotic duration observed in HeLa cells (Supplementary Figure S2C). We also did not observe defects in chromosome alignment or segregation (Supplementary Figure S2D). Thus, the kinetochore-localized fraction of the phosphodomain does not detectably perturb chromosome biorientation or cell-cycle progression.

### Bub1 and BubR1 recruited by the KI motifs do not contribute to MCC assembly mediated by the MELT motifs

We first determined the baseline activity of the four MELT motifs by establishing the dose-response correlation for the eSAC phosphodomain with inactive KI motifs (KI1*-KI2* in Figure 2B left, purple circles). The response elicited by this eSAC phosphodomain was non-monotonic: the mitotic duration increased steeply before gradually decaying to a lower value (see Supplementary Figure S3A). A non-monotonic response was not apparent for a previously characterized eSAC phosphodomain containing four MELT motifs (numbers 11-14) (Chen et al., 2019). The different behaviors of the two phosphodomains may be ascribed to different Bub1-Bub3 binding affinities of the MELT motifs that they contain. Notably, when both KI motifs were active (KI1-KI2), the maximal duration of mitosis was significantly reduced (∼ 87 minutes estimated by 4-parameter sigmoidal fit to the binned averages of the data, see Figure 2C). The KI motifs similarly suppressed the signaling strength of an extended phosphodomain containing 7 MELT motifs (Supplementary Figure S3B).

We next determined the response elicited by the two eSAC phosphodomains containing four MELT motifs and only one active KI motif. When only the first KI motif was active (KI1-KI2*, only Bub1 depleted, see Figure 2B), the dose-response data were monotonic with a slightly lower maximal response than the maximal response elicited by KI1*-KI2* (Figure 2C green circles and curve). When only the second KI motif was active (KI1*-KI2, Bub1, and BubR1 depleted, Figure 2C), the maximal response was significantly attenuated (blue circles and line in Figure 2C). Interestingly, the response to this eSAC was also non-monotonic, an initial overshoot followed by decay to a lower response level (residuals from the Hill equation fit shown in Supplementary Figure S3A).

These results reinforce the conclusion that Bub1 and BubR1 recruited by the two KI motifs do not directly contribute to MCC generation by the MELT motifs within the same phosphodomain (Figure 2C). The distinctly different effects of Bub1 and BubR1 sequestration on eSAC activity also suggest that the cellular abundance of these proteins may be an important aspect of SAC signaling.

### Numerical simulation of the dose-response data

The strong suppression of eSAC signaling by the second KI motif that binds BubR1 can be ascribed to two effects of BubR1 sequestration: reduced rate of MCC assembly and a lowered limit on the maximal amount of MCC that can be generated. The latter effect is unlikely to play a major role in shaping the observed dose-response data. This is because the KI1*-KI2* eSAC delays mitosis by at most 300 minutes, significantly shorter than ∼ 1500 minutes long arrest seen in nocodazole-treated HeLa cells (Collin et al., 2013; Dick and Gerlich, 2013). Therefore, the amount of MCC produced by the eSAC systems is likely to be lower than the amount produced in nocodazole-treated HeLa cells. Therefore, the effect of BubR1 sequestration on the *rate of MCC generation rather than the maximal amount of MCC that can be generated* likely shapes the dose-response data.

An intuitive explanation for the dose-response dependence can be developed using the following four observations: (1) Bub1 and BubR1 interactions with the KI motifs do not require MELT motif phosphorylation (Figure 2B), (2) Bub1 and BubR1 recruited by the KI motifs do not contribute to the activity of the phosphorylated MELT motifs (Figure 2A-C), (3) the eSAC phosphodomains are significantly more abundant than Bub1 and BubR1 (shown later), and (4) phosphorylated MELT motifs in an eSAC phosphodomain recruit Bub1, BubR1, and Mad1 (Chen et al., 2019). The first three observations indicate that the two KI motifs in the eSAC phosphodomains will differentially sequester Bub1 or BubR1, and the last observation suggests that the MELT motifs in the eSAC phosphodomains will form two distinct signaling complexes: MELpT:Bub1:Mad1/2 and MELpT:Bub1:BubR1:Mad1/2 (Figure 3A; Cdc20 and Mad1/2 are present in both and hence not indicated). When taken together, these observations suggest that the differential sequestration of Bub1 and BubR1 by the KI motifs will affect the composition of the signaling complexes assembled on the phosphodomains. If the two signaling complexes assemble the MCC at different rates, the result will be different mitotic delays explaining why the four phosphodomains produce distinctly different maximal mitotic delays (Dick and Gerlich, 2013). The quantitative dose-response data and numerical simulations provide an excellent opportunity to test this model and the notion that MELpT:Bub1:BubR1:Mad1/2 generates MCC at a higher rate than the MELpT:Bub1:Mad1/2.

**Figure 3.**
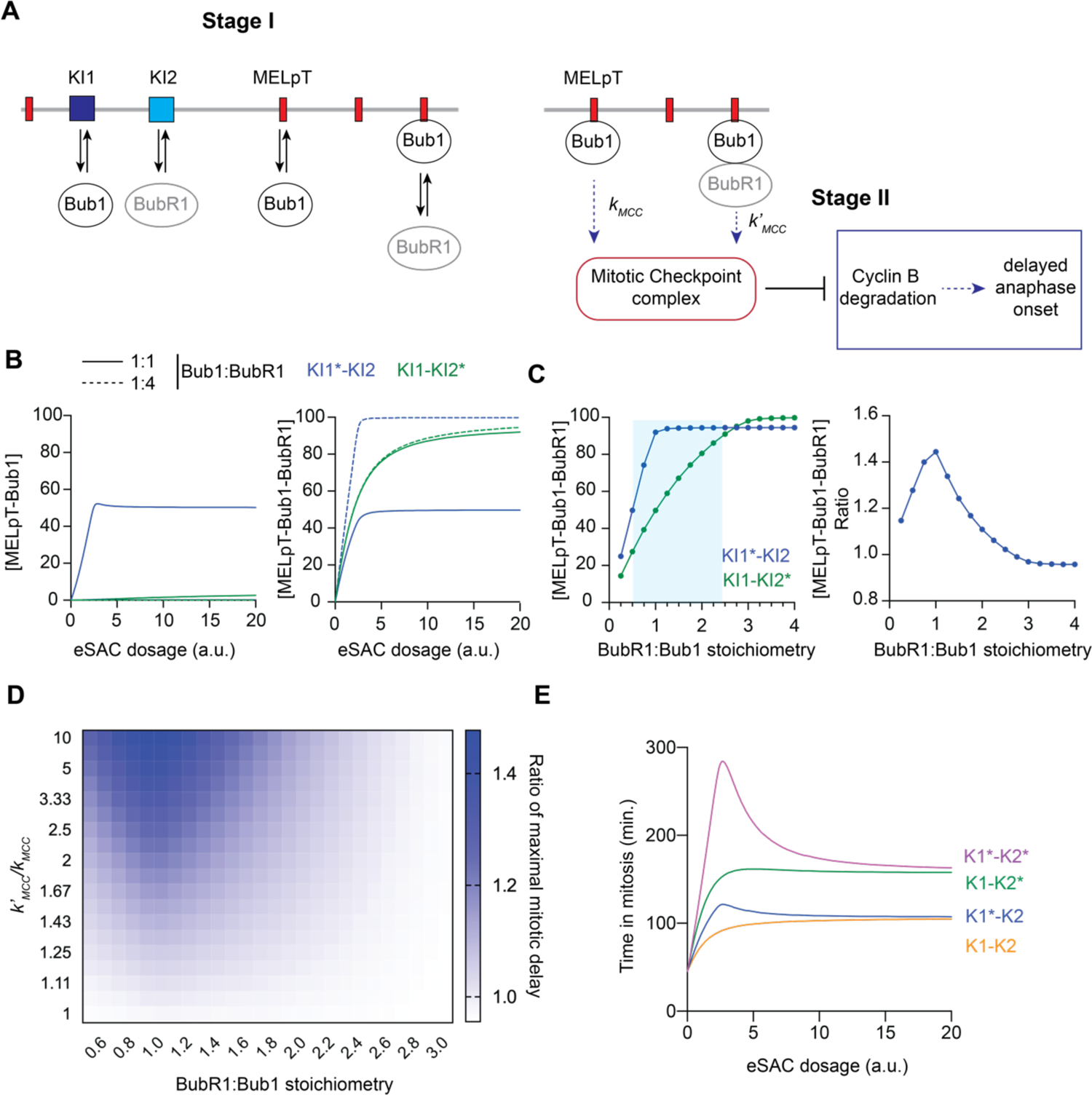
Numerical simulation of the dose-response data for the eSAC phosphodomains containing KI motifs. **A** Schematic of the two-stage model used to simulate the dose-response curves. **B** The influence of Bub1-BubR1 stoichiometry and the differential sequestration of Bub1 and BubR1 by the KI motifs on the equilibrium concentrations of the two signaling complexes formed on phosphorylated MELT motifs. **C** Comparison of the equilibrium concentration of the MELpT-Bub1-BubR1 complex assembled by KI1*-KI2 and KI1-KI2* (left) and the ratio (right) of their maximal responses as a function of the Bub1-BubR1 stoichiometry. For the two phosphodomains to generate different responses, the Bub1-BubR1 stoichiometry must be around 1:1 (indicated by the blue shaded area). **D** The ratio of the maximal responses produced by KI1*-KI2 and KI1-KI2* as a function of the Bub1-BubR1 stoichiometry and the ratio of the rates at which the MELpT-Bub1-BubR1 and the MELpT-Bub1 signaling complexes produce MCC (i.e., *k’_MCC_/k_MCC_*). **E** Simulation of the dose-response curves for the four phosphodomains using the same set of parameter values.

We simulated this model in two stages (Figure 3A). In the first stage, we calculate the equilibrium concentrations of the two signaling complexes using rates governed by mass action (equations 6-15, Methods); in the second stage, we calculate the rate of MCC formation and its effect on the metaphase-to-anaphase transition (equations 16-17, Methods). For both signaling complexes, the events leading up to Mad2:Cdc20 formation are identical. Therefore, we did not explicitly simulate them. After this step, MELpT:Bub1:BubR1 can assemble the MCC either within the signaling complex itself or in the cytosol, whereas MELpT:Bub1 must rely on cytoplasmic MCC assembly. Using these insights, we expanded our previously described eSAC model (Chen et al., 2019). This model simulates the activity of the four MELT motifs in a manner analogous to the original model. Bub1-Bub3 and BubR1-Bub3 complexes are represented by ‘Bub1’ and ‘BubR1’ (Figure 3A) (Overlack et al., 2017; Overlack et al., 2015). Activities of the two KI motifs are simulated as protein-protein interactions assuming that KI1 interacts only with Bub1, and KI2 interacts with only BubR1 (Figure 3A). Finally, the KI motifs and phosphorylated MELT motifs are assumed to act independently.

### Dependence of the equilibrium concentrations of MELpT:Bub1 and MELpT:Bub1:BubR1 assembled by the eSAC systems on Bub1-BubR1 stoichiometry

The equilibrium concentrations of MELpT:Bub1 and MELpT:Bub1:BubR1 depend on two sets of parameters: (1) the concentrations of Bub1, BubR1, and the eSAC phosphodomain (quantification shown after the simulations), (2) the affinities of Bub1 and BubR1 for KI1 and KI2 respectively, Bub1 for phosphorylated MELT motifs, and BubR1 for Bub1 (Krenn et al., 2014; Vleugel et al., 2015; Zhang et al., 2016). We assume that Bub1-BubR1 heterodimerization occurs after Bub1 binds MELpT. However, this assumption will not affect the behavior of the model. The values of the parameters used are listed in Table S2.

Our first goal was to delineate the effects of the two KI motifs by understanding the reason for the different mitotic durations achieved by KI1*-KI2 and KI1-KI2* at high eSAC dosages. For these phosphodomains to elicit different maximal responses, they must form different amounts of MELpT:Bub1 and MELpT:Bub1:BubR1. Therefore, we investigated how the above two parameter sets affect the equilibrium concentrations of the two signaling complexes formed by KI1*-KI2 and KI1-KI2*. Given a set of affinities of KI1 and KI2 for Bub1 and BubR1, respectively, the concentrations of MELpT:Bub1 and MELpT:Bub1:BubR1 will depend on the relative cellular abundance of Bub1 and BubR1. If BubR1 is much more abundant than Bub1, then nearly every MELpT:Bub1 signaling complex will be able to recruit BubR1 to form MELpT:Bub1:BubR1. Consequently, the concentration of MELpT:Bub1 will become negligible (dashed green and blue lines respectively near the X-axis in Figure 3B, left). At high eSAC dosage, both eSAC systems will form similar amounts of MELpT:Bub1:BubR1 despite the differential sequestration of Bub1 and BubR1 (converging dashed lines at high eSAC dosage in Figure 3B, right). Therefore, their eSAC signaling activity will also be similar, contrary to our observations.

When BubR1 and Bub1 concentrations are similar, a sizeable fraction of MELpT:Bub1 will not recruit BubR1 (Figure 3B, right, solid lines) especially at a high eSAC dosage. Therefore, at high eSAC dosage, the concentration of MELpT:Bub1:BubR1 is lower for KI1*-KI2 than for KI1-KI2* (solid lines in Figure 3B, left). Figures 3C shows how Bub1-BubR1 stoichiometry affects MELpT:Bub1:BubR1 concentration for the high and constant eSAC dosage (Figure 3C left). BubR1:Bub1 < 2.5 ensures that the two phosphodomains will assembly different concentrations of MELpT:Bub1 and MELpT:Bub1:BubR1 (blue shaded region in Figure 3C left). This difference in the equilibrium concentrations of the signaling complexes can form the basis for differential eSAC activities.

### A higher rate of MCC generation by MELpT:Bub1:BubR1 compared to MELpT:Bub1 can explain the differential behavior of the two phosphodomains

Differences in the concentrations of MELpT:Bub1 and MELpT:Bub1:BubR1 will translate into different activity only if they generate MCC at different rates. As before, we calculate the rate of MCC generation by assuming it to be proportional to the concentrations of MELpT:Bub1 and MELpT:Bub1:BubR1 (Figure 3B). Following our experimental results, the KI-bound Bub1 and BubR1 do not promote eSAC signaling. To simulate the effect of the MCC generated on mitotic progression, we modified the model of metaphase-to-anaphase transition described by He et al. (Figure S4B, Methods) (Chen et al., 2019; He et al., 2011).

As discussed in the preceding sections, the maximal mitotic duration produced by the two eSAC systems depends on: (1) the equilibrium concentrations of the MELpT:Bub1 and MELpT:Bub1:BubR1, which are in turn dependent on the Bub1-BubR1 stoichiometry, and (2) the values of the MCC generation rate constants *k_MCC_* and *k’_MCC_*. To determine the working combination of these factors, we fixed the eSAC dosage at a high value and calculated the duration of mitosis for a range of ratios of *k_MCC_* to *k’_MCC_* and Bub1:BubR1 stoichiometry. Figure 3D displays how the two ratios affect the maximal duration of mitosis achieved by KI1-KI2* and KI1*-KI2, respectively. In particular, for Bub1:BubR1 ∼ 1 and *k_MCC_/ k’_MCC_* > 6, the ratio of the maximal mitotic durations achieved by the KI1-KI2* and KI1*-KI2 eSAC systems exceed 1.4.

Following this result, we used [Bub1]:[BubR1] = 1 and *k_MCC_:k’_MCC_ =* 0.1 (rate constant for MCC assembly within the signaling complex is 10-fold higher than the rate constant for cytoplasmic MCC assembly) to simulate the dose-response curves for all four phosphodomains. This simulation captures key characteristics of the dose-response data for all four phosphodomains (Figure 3E). As before, the assumption of synergistic signaling was necessary to reproduce the non-monotonicity of the dose-response data for the phosphodomain containing four MELT motifs (Chen et al., 2019). Without it, the responses elicited by all phosphodomains become monotonic (Figure S4D).

### Stoichiometry of Bub1, BubR1, Mad1, and the eSAC phosphodomain in HeLa cells

The relative amounts of Bub1, BubR1, and the eSAC phosphodomain emerge as critical determinants of eSAC signaling. Therefore, for comparative protein abundance measurements, we constructed genome-edited HeLa cell lines wherein mNeonGreen (abbreviated as mNG) was fused to the N-terminus of Bub1 and BubR1 and the C-terminus of Mad1. In all three cases, we obtained heterozygous cell lines with partially edited genomes. Consequently, approximately half of the protein in these cells was labeled (Figure 4A, Methods).

**Figure 4.**
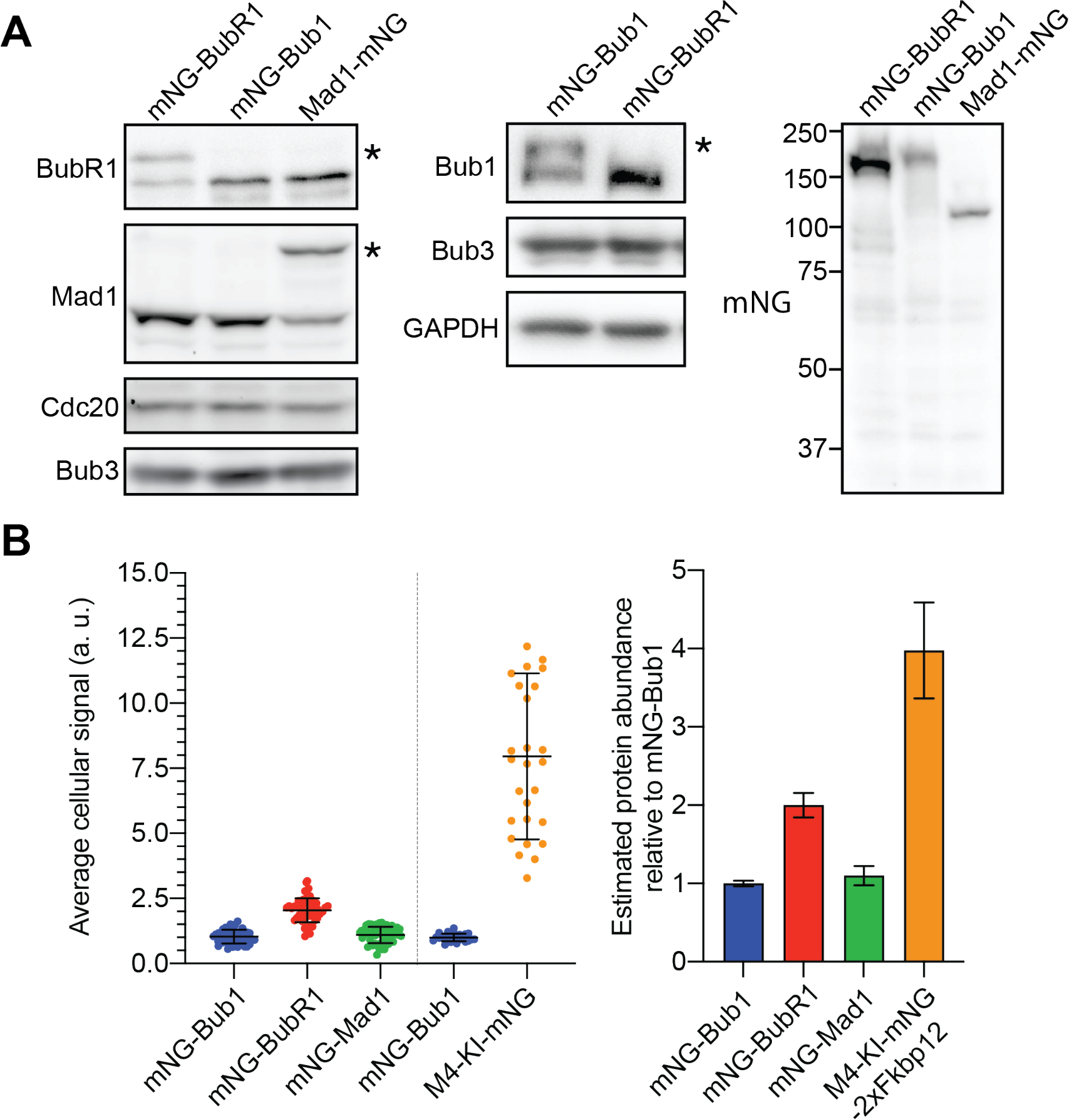
Quantification of the relative abundance of a eSAC phosphodomain, mNeonGreen-Bub1, mNeonGreen-BubR1, and Mad1-mNeonGreen in genome-edited HeLa cell lines. **A** Immunoblots showing that roughly half of BubR1 and Mad1 (left) and Bub1 (middle) proteins in the three partially genome-edited cell lines is tagged with mNeonGreen (asterisks on the right of each displayed blot mark the mNG fusion protein). Immunoblot on the right shows whole-cell extracts of the three cell lines probed with anti-mNeonGreen antibodies. **B** Average mNG-Bub1, mNG-BubR1, Mad1-mNG, and M4-KI1-KI2-mNG-2xFkbp12 signals from mitotic HeLa cells (left) and estimation of the relative protein abundance, assuming that the total protein abundance is twice as high as the abundance of the mNG-labeled species (mean ± std. dev.).

Quantitation of mNG fluorescence in mitotic cells revealed that the KI1-KI2 phosphodomain was 2-4-fold more abundant than the three SAC proteins (Figure 4B, Methods). Therefore, and given the constitutive activity of the two KI motifs, the eSAC phosphodomains containing active KI motifs will deplete Bub1 and BubR1 from the cytosol. The measurements also show that BubR1 is ∼ 2-fold more abundant than Bub1 and Mad1 (Figure 4B). In the case of BubR1, the measured abundance includes free BubR1-Bub3 and BubR1-Bub3 incorporated into MCC. Although the fraction of free BubR1-Bub3 remains unknown, this value is likely to be less than 2-fold higher than that of Bub1-Bub3, consistent with the requirement of comparable amounts of Bub1-Bub3 and BubR1-Bub3 in our simulations.

### Recruitment of BubR1 by Bub1 per se strengthens kinetochore-based SAC signaling

Following these results and insights, we re-examined the role of Bub1-BubR1 heterodimerization in kinetochore-based SAC signaling. PP2A recruitment to the kinetochore in this manner contributes to SAC silencing directly (Espert et al., 2014; Kruse et al., 2013; Qian et al., 2017), and indirectly by either promoting PP1 recruitment (Nijenhuis et al., 2014) or stabilizing kinetochore-microtubule attachment (Suijkerbuijk et al., 2012). Therefore, we created two BubR1 mutants: mNG-BubR1^ΔKARD^ that lacks the PP2A-binding KARD domain and mNG-BubR1^ΔHD, ΔKARD^ that additionally lacks the BubR1 heterodimerization domain (Figure 5A). The complete removal of KARD in these mutants is expected to abolish the binding of B56α and other isoforms of B56 onto BUBR1 (Wang et al., 2016a; Wang et al., 2016b) enabling us to analyze whether the recruitment of BUBR1 to the signaling kinetochore *per se* contributes to the SAC activity. Importantly, we ensured that the expression level of these mutants was similar to wildtype BubR1 because the transient over-expression of BubR1 can deplete the cytosolic pool of Bub3 (Taylor et al., 1998) and adversely affect the SAC or even induce cell death (data not shown). Higher cytosolic BubR1 concentration may also proportionally increase the rate of cytosolic MCC formation and thus mask impaired MCC assembly within the kinetochore.

**Figure 5.**
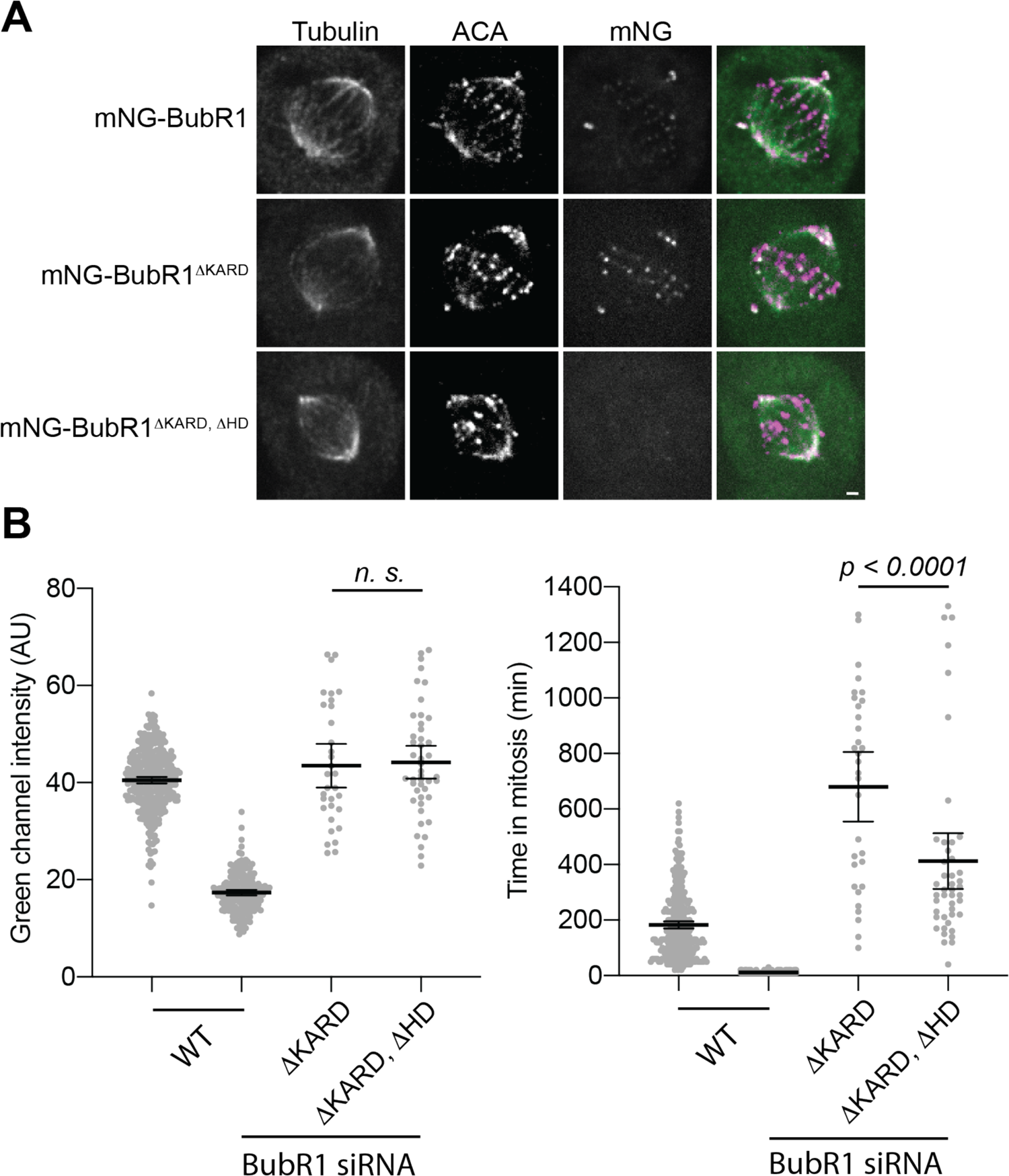
Recruitment of BubR1 by Bub1 per se contributes to the activity of the kinetochore-based SAC signaling. **A** Micrographs of representative cells displaying the indicated antigens at the top. The top row displays micrographs of the partially genome-edited cell line expressing mNG-BubR1. The bottom two rows display representative cells after endogenous BubR1 was knocked down and the indicated mutant, mNG-tagged version was ectopically expressed (scale bar ∼ 1.22 microns). **B** Scatterplot on the left displays the duration of mitosis of either genome-edited *mNG-BUBR1* or cells that ectopically expressed the indicated mutant after treatment with BubR1 siRNA in media containing 25 nM nocodazole. Scatterplot on the right displays the average cytosolic mNeonGreen signal from the same cells (mean ± 95% confidence intervals).

We knocked down endogenous BubR1 in HeLa-A12 using RNA interference and rescued these cells with mNG-BubR1^ΔKARD^ or mNG-BubR1^ΔHD, ΔKARD^. As expected, mNG-BubR1 ^ΔKARD^ localized to unattached kinetochores, whereas mNG-BubR1^ΔHD, ΔKARD^ localization to unaligned kinetochores was undetectable (Figure 5A). We also quantified the duration of mitotic arrest using nocodazole treatment following the previous studies. To ensure that the BubR1 mutants were not over-expressed, we established the physiological BubR1expression level by quantifying cytosolic BubR1 fluorescence in the genome-edited *mNG-BUBR1* HeLa-A12 cells treated with control siRNA and imaged under identical conditions. In our knock-in/knock-down experiments, we considered only those cells exhibiting mNG intensity that is 0.5–2-times the average mNG intensity of mitotic mNG-BubR1 HeLa-A12 cells (Figure 5B, left). Quantification of mitotic duration revealed that cells expressing mNG-BubR1^ΔKARD^ arrested significantly longer than control cells; the longer duration is attributable to the loss of PP2A activity from the kinetochores (Saurin et al., 2011). Notably, cells rescued with mNG-BubR1^ΔHD, ΔKARD^ arrested for a significantly shorter amount of time compared to the cells rescued with mNG-BubR1^ΔKARD^ (Figure 5B, right). Thus, the recruitment of BubR1 *per se* strengthens the SAC.

In conclusion, Bub1-BubR1 heterodimerization significantly enhances SAC signaling activity in human cells. This enhancement may be simply due to enrichment of BubR1 at the site of formation of Mad2-Cdc20 (Lara-Gonzalez et al., 2021a; Piano et al., 2021), although more complicated mechanisms can also be envisioned. Bub1 and BubR1 recruitment via the KI motifs does not contribute to eSAC signaling mediated by MELT motifs indicating that their contribution to SAC signaling is likely to be minor. Although prior studies found that the KI motifs promote SAC signaling, this contribution was detectable only in the context of recombinant Knl1 variants containing either just one MELT motif (Krenn et al., 2014) or three inactive MELT motifs (Vleugel et al., 2013); the contribution was undetectable in a Knl1 variant containing multiple MELT motifs but lacking the N-terminus including the KI motifs (Zhang et al., 2014). Compared to the KI motifs, phosphorylated MELT motifs bind significantly more strongly to Bub1 and BubR1 (Krenn et al., 2012; Primorac et al., 2013; Zhang et al., 2016). MELT motifs also vastly outnumber the KI motifs. Therefore, phosphorylated MELT motifs recruit the majority of Bub1 and BubR1 to the kinetochore (Vleugel et al., 2015; Zhang et al., 2014). Our findings highlight the dual effect of Bub1-mediated BubR1 recruitment on the SAC. BubR1 stabilizes the kinetochore-microtubule attachment by recruiting PP2A, thereby promotes the silencing of SAC. But it also promotes the SAC activity per se, which is critical for minimizing chromosome missegregation in normally dividing cells, wherein the last few unattached kinetochores need to be able to signal the cell to delay anaphase onset (Roy et al., 2020).

## Acknowledgements

This work was supported by R35-GM126983 from NIGMS to APJ. We thank Dr. Stephen S. Taylor for the generous gift of antibodies, Dr. Eugene Makeyev for the HeLa acceptor cell line, and Dr. Michael Lampson for plasmids. We also thank members of the Joglekar lab for stimulating discussions. The use of the Incucyte system was made possible by a generous gift from the Richard Tam Foundation to Prof. Sue O’Shea (Cell & Developmental Biology, University of Michigan Medical School).

**MATLAB codes used for the simulations and for generating figure panels can be accessed on GitHub:** https://github.com/anandban/eSAC-KI)

**The authors declare that no competing financial interests exist.**

## Materials and Methods

### Plasmid construction

The plasmids used for the stable cell lines were based on plasmids that have been described previously (Chen et al., 2019). Briefly, the phosphodomain was integrated into either *Not*I or *Asc*I and *Xho*I restriction sites to create constitutively expressed phosphodomain-mNeonGreen-2xFkbp12. The Mps1^500-857^ fragment corresponding to the Mps1 kinase domain was integrated into the *Fse*I and *Bgl*II restriction sites to create conditionally expressed Frb-mCherry-Mps1^500-857^. eSAC phosphodomains spanning the 1^st^ MELT motif (M1) and the two KI motifs were created by fusing Knl1^160-256^ to mNeonGreen-2xFkbp12. eSAC phosphodomains containing four MELT motifs and the two KI motifs were created by fusing Knl1^160-256^ to Knl1 fragment Knl1^881-1014^ spanning MELT motifs 12-14, which has been characterized previously (Vleugel et al., 2015). The activity of the first KI motif was disrupted by changing its amino acid sequence from ‘K**I**D**TT**S**F**LANLK’ to ‘KADAASALANLK’ (KI1*). Similarly, the activity of the second KI motif was disrupted by mutating its amino acid sequence from ‘KIDFND**F**IK**R**LK’ to ‘KIDFNDAIKALK’ (KI2*) following (Krenn et al., 2012).

DNA repair templates used for CRISPR/Cas9 mediated genome editing were constructed via DNA assembly using the NEB HiFi DNA assembly kit per the manufacturer’s instructions. Successfully edited alleles encode mNeonGreen-tagged SAC proteins that separate the corresponding wildtype protein and the fluorescent protein mNeonGreen by a short flexible linker (mNG-BUBR1 and mNG-BUB1: GSGGSG; MAD1-mNG: GGAGGSGG). The sequences of all homology-directed repair template plasmids and Cre-lox recombination-mediated cassette exchange plasmids are available upon request.

### Tissue culture and cell line construction for eSAC analyses

Henrietta Lacks (HeLa) cells were grown in DMEM media supplemented with 10% FBS, 1% Pen/Strep, 1x-Glutamax, and 25 mM HEPES under standard tissue culture conditions (37 °C and 5% CO2). Stable cell lines expressing the two eSAC components were generated by integrating a bi-cistronic eSAC plasmid at engineered *lox* sites in the HeLa genome according to the protocol described in (Khandelia et al., 2011). Upon transfection, DMEM media supplemented with 1 μg/ml Puromycin was used to select transformed cells, and all the colonies were pooled to culture the transformed cells used in the experiments.

To conduct dose-response analysis, each eSAC cell line was plated ∼ 40-48 hours prior to the start of the experiment in DMEM media without Puromycin. Doxycycline was added at the time of plating to induce the expression of Frb-mCherry-Mps1. Prior to imaging, the cells were washed with PBS. Fluorobrite media with 10% FBS 1% Pen/Strep with or without Rapamycin were added to each well.

### Genome editing HeLa cells using CRISPR/Cas9

The guide RNAs (gRNAs) for in situ BUBR1 and BUB1 N-terminal mNeonGreen-tagging were 5’-CAGGAUGGCGGCGGUGAAGA-3’ and 5’-GGUUCAGGUUUGGCCGCUGC-3’, respectively. The gRNA for in situ MAD1 C-terminal mNeonGreen-tagging was 5’-CAGACCGUGGCGUAGCCUGC-3’. Single guide RNAs (sgRNAs) were synthesized using the EnGen sgRNA Synthesis Kit (for the *Streptococcus pyogenes*-originated Cas9, New England Biolabs). The SpCas9-sgRNA ribonucleoprotein (RNP) complex was assembled at room temperature in a buffer consisting of 20 mM of HEPES-KOH (pH 7.5), 150 mM of KCl, 1 mM of MgCl_2_, ten percent (by volume) of glycerol, and 1 mM of DTT using 100 pmol of SpCas9-2×NLS (the QB3 MacroLab) and 120 pmol of sgRNA. The RNP complex and 1.5 μg of a linearized homology-directed repair template plasmid were transfected into 2×10^5^ – 5×10^5^ nocodazole-arrested mitotic HeLa A12 cells using a Nucleofector and the associated Cell Line Kit R (Lonza) following manufacturer’s instructions. After five weeks, green fluorescence-positive mitotic cells (arrested by 330 nM of nocodazole for 16 hours) were sorted directly into 96-well plates at 1 cell/well. Healthy colonies were subject to further validation by genotyping and sequencing, as well as immunoblotting.

For genotyping, HeLa-A12 genomic DNAs were purified using the Wizard SV Genomic DNA Purification System (Promega). Genotyping primers (BUBR1 forward primer 5’-CCTGGTCACATCTGAGCTAT-3’, BUBR1 reverse primer 5’-CTCAGTGAGACTCCAGTGTT-3’, BUB1 forward primer 5’-CCCTCTACATGAAGGCGCTA-3’, BUB1 reverse primer 5’-GCTCGCCCAAGGTAAACATT-3’, MAD1 forward primer 5’-GGACTTTTCAGGGACGTGGT-3’, and MAD1 reverse primer 5’-GAGTTGGGAGGAGGGGACTC-3’) were designed to bind outside of homology arms to avoid false-positive colonies from integration of the homology-directed repair template plasmid to an off-target genomic locus.

### Drug and RNAi treatments

To induce the expression of the mCherry-Frb-Mps1 kinase domain doxycycline was added to a final concentration of 2 ug/ml (stock concentration 2 mg/ml in DMSO) ∼ 48 hours prior to the start of the experiment. Prior to prior to the start of each experiment, Rapamycin was added ∼ 1 hour of the experiment to a final concentration of 500 nM (stock concentration 500 μM in DMSO) to induce the dimerization of the eSAC kinase domain with the eSAC phosphodomain. Nocodazole was added to the final concentration of 330 nM (stock concentration 330 μM in DMSO). The cocktail of siRNA against five different B56 isoforms was added to a final concentration of 40 nM (stock concentration 10 μM). The siRNA sequences were obtained from ref. (Nijenhuis et al., 2014). Cell cycle synchronization was in G1/S was achieved by treating cells with 2.5 mM thymidine (from a 100 mM stock in PBS) for 16-18 hours. Cells were washed with DMEM for release from the G1/S arrest. To arrest cells in a prometaphase-like condition, cells were released from a G1/S block and then treated with 236 nM GSK923295 (stock concentration 236 μM in DMSO) ∼ seven hours post-release and imaged after one hour.

### Immunoprecipitation

HeLa A12 cells constitutively expressing either MELT1-KI or MELT1-KI-M3 were synchronized at G1/S by 2.5 mM of thymidine. 9 hours after being released from the (double) thymidine block, cells were synchronized in metaphase using 10 μM MG132. After another 1.5 hours, cells were scraped off the plate, washed once with PBS, pelleted, snap-frozen, and stored at −80 °C. Cells were thawed, resuspended in the complete lysis buffer (75 mM HEPES-HCl (pH 7.5), 150 mM KCl, 1.5 mM EGTA, 1.5 mM MgCl2, 10% (by volume) glycerol, and 0.075% (by volume) Nonidet P-40 (AmericanBio); immediately before use one cOmplete protease inhibitor cocktail tablet (EDTA-free, Roche Diagnostics) and a phosphatase inhibitor cocktail (1 mM Na_4_P_2_O_7_, 0.1 mM Na_3_VO_4_, 5 mM NaF, and 2 mM sodium β-glycerophosphate) were added), and lysed for 1 hour at 4 °C while rotating. Cell lysates were then centrifuged at 18,000 g for 25 min at 4 °C. The supernatant was subsequently cleared by control agarose beads for 1.5 hours at 4 °C to reduce non-specific binding. Cleared supernatant was then mixed with mNeonGreen-Trap agarose beads (ChromoTek) and rotated for 1.5 hours at 4 °C. Beads were washed 4 times using the lysis buffer. Finally, 2×Laemmli buffer (Bio-Rad Laboratories) supplemented with β-mercaptoethanol was added to the beads. Samples were boiled in a water bath for 10 min before subjecting to SDS-PAGE. The following antibodies and working dilutions were used in immunoblotting:

SB1.3 antibody against BUB1 (Taylor et al., 2001), 1:500 or 1:1000, sheep polyclonal; BUB3 antibody (Sigma B7811), 1:500, rabbit polyclonal; FKBP12 antibody (Abcam ab2918), 1:2000, rabbit polyclonal; BUBR1 antibody (Bethyl A300-995A), 1:1000, rabbit polyclonal.

### Immunofluorescence

HeLa cell lines were grown on sterile coverslips in 6 well plates in media supplemented with 1 μg/ml Doxycycline to induce the expression of mCherry-Frb-Mps1 kinase domain. After ∼ 48 hours, the cells were treated with 500 nM Rapamycin. After four hours of incubation, cells were fixed with 4% paraformaldehyde and stained with ACA antibodies (1:1000, Antibodies Inc., Davis, CA) and Alexa-633-conjugated secondaries (1:5000). After staining, the cells were embedded in Diamond mounting media and stored at room temperature.

The mounted cells were imaged on an Eclipse Ti-E/B inverted microscope (Nikon) with a CFI Plan Apochromat VC 100 ×, 1.40 NA oil objective (Nikon). The microscope was equipped with a H117E1 motorized stage (Prior Scientific), a NanoScanZ 100 piezo stage (Prior Scientific), and an X-Light V2 L-FOV confocal unit with 60 μm pinholes (CrestOptics). A CELESTA Light Engine (Lumencor) served as the excitation laser source, featuring a 477-nm line for imaging the mNeonGreen protein, 546-nm line for imaging mCherry, and 647-nm line for imaging the Alexa-633-conjugated antibodies. Fluorescence emission light was filtered by ET605/52m (Chroma Technology) for the red channel and by ET525/36m (Chroma Technology) for the green channel. Images were acquired by a Prime 95B 25mm sCMOS camera (Teledyne Photometrics). A custom MATLAB program was used to quantify kinetochore-localized fluorescence signals.

### Long term live-cell imaging of HeLa cells

Imaging was conducted as described in detail previously (Chen et al., 2019). We used either the Incucyte Zoom live cell imaging system (Sartorus) or the ImageXpress Nano live cell imaging system (Molecular Devices), both equipped with a 20x Phase objective. To image cells on the Incucyte system, cells were plated in 12-well plastic tissue culture plates, whereas they were plated in 24-well plate glass-bottom dishes (Corning) for the ImageExpress Nano system. At each position, one phase, GFP, and mCherry image was acquired every 10 minutes. The exposure times for mCherry and GFP images were adjusted to minimize photobleaching while enabling accurate determination of intensity values. It should be noted that the excitation sources, optics, and detector on the ImageXpress Nano and the Incucyte microscope are entirely different. Therefore, the mCherry intensity values across different experiments are not directly comparable.

### Image analysis

Prior to intensity quantification, acquired images were pre-processed using functions from the ‘Image Processing Toolbox’ provided with Matlab as follows. First, the phase image sequence was registered to remove any movement of the field of view between adjacent timepoints, and at each timepoint, the same transform was applied to the GFP and mCherry images to register them.

Additionally, image intensity from a blank, unseeded well was used for background correction of the fluorescence channels. Next, GFP and mCherry fluorescence signals were quantified using a custom graphical user interface (GUI) written in Matlab as described previously. Briefly, this interface uses cross-correlation of each phase image with a circular kernel to identify cells with circular shapes close to the diameter of the circular kernel. The centroids of these shapes were then linked along the time axis. These images were presented to the user via the GUI to (a) discard false positive, non-mitotic cells or debris and (b) visually correct the time of entry into or exit from mitosis. The GUI then calculated the GFP and mCherry signals per cell as the average fluorescence intensity.

In all the dose-response assays discussed in this study, the phosphodomain is highly and constitutively, expressed in all cells, whereas the kinase domain is expressed conditionally by an inducible promoter. Consequently, the amount of the kinase domain expressed varies from cell to cell, and it is lower than the amount of the phosphodomain in most cells. Because of this design of the eSAC system, the dosage of the dimerized signaling complex in a cell can be inferred from the amount of Frb-mCherry-Mps1^500-817^ in that cell. Therefore, we defined the dosage of the eSAC signaling complex by quantifying the average mCherry fluorescence within a cell and the response as the duration of mitosis (the amount of time that the cell spends with a spherical morphology that is characteristic of mitotic HeLa cells).

The Hill equation is used to fit the sigmoidal trend:

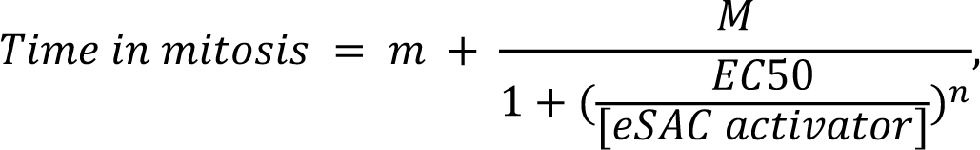

wherein *n* is the Hill coefficient and *EC50* is the level of the eSAC activator at which the time in mitosis reaches the middle between the baseline level (*m*) and the plateau level (*m* + *M*).

### Statistical analysis

To determine the overall trend in the dose-response data, the data were first binned (in MATLAB), and then the mean values of each bin were overlayed on the data. The number of observations and technical replicates are noted in the figure legends. These mean values were fit with a four-parameter sigmoidal curve using the GraphPad Prism 9 software. Statistical significance of the difference between the mean values in Figure 5B was assessed using the unpaired t-test with Welch’s correction.

## Mathematical modeling

### Modeling the activity of Bub1 phosphodomains

#### Stage 1: Calculation of the steady-state concentrations of signaling complexes

(MATLAB codes available on GitHub: https://github.com/anandban/eSAC-KI)

This model simulates the eSAC activity of the Bub1^221-620^ and Bub1^441-620^ phosphodomains. In the equations below, we refer to these phosphodomains simply as ‘Bub1’. The eSAC activator complex is formed by the dimerization of Bub1 with the Mps1 kinase domain (Bub1:Mps1). Once Mps1 phosphorylates Bub1, Bub1 can bind Mad1/2 (Bub1:Mps1:Mad1/2). Therefore, the concentrations of different species of Bub1 are related by the equation:

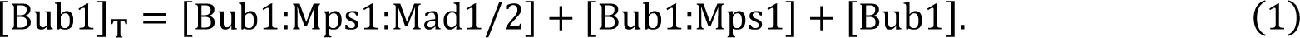

Assuming reversible binding between phosphorylated Bub1 and Mad1/2 complex, the concentration of eSAC activator complex will saturate to a value dependent on the finite concentration of Mad1/2 (set at 100 nM, Fig. S1A). Note that Bub1 can produce MCC only if it recruits Mad1/2. Therefore, even though Bub1:Mps1 and Bub1 can both bind BubR1 and Cdc20, they do not participate in SAC signaling. The recruitment of SAC proteins, formation of signaling complexes, and MCC are calculated by assuming mass action kinetics.

#### Model of Bub1-mediated MCC formation

Available data suggest that a signaling complex comprising Bub1^221-620^, BubR1, Mad1, and Cdc20 facilitates the formation of either Mad2:Cdc20 or the MCC. We assume that when BubR1 is present the phosphodomain assembles MCC, and when BubR1 is absent, the phosphodomain produces Mad2:Cdc20. Therefore, the rate of Mad2:Cdc20 and MCC formation is calculated as:

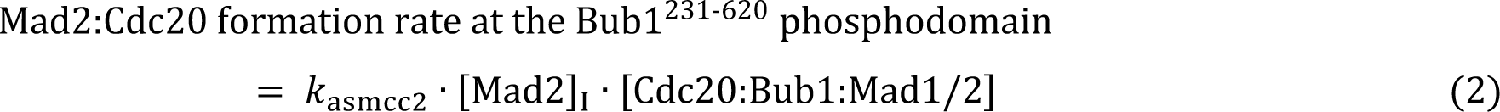

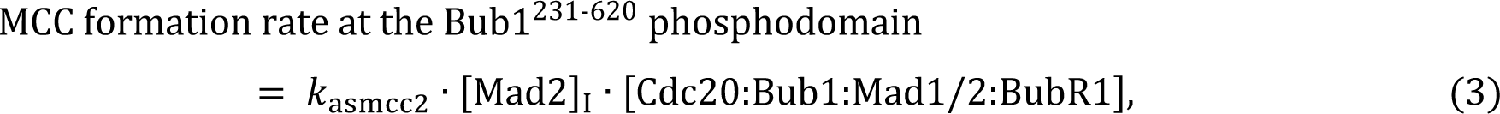

where, [Mad2]_I_ = the concentration of inactive (open) form of Mad2 in the cytoplasm [Cdc20:Bub1:Mad1/2] = the concentration of the complex between Bub1, Mad1, and Cdc20. [Cdc20:Bub1:Mad1/2:BubR1] = the concentration of the complex between Bub1, BubR1, Mad1, and Cdc20.

For the sake of simplicity, we assume that the rate constant for MCC formation (*k_asmcc2_*) is numerically equal to the rate constant for Mad2:Cdc20 formation. This assumption is consistent with the observation that Mad2:Cdc20 formation is the rate-limiting step in MCC formation (Faesen et al., 2017).

The cytosolic Mad2:Cdc20 molecules produced by either phosphodomain interact with cytosolic, free BubR1 to complete MCC formation. We denote the rate constant for this reaction by *k_asmcc1_*. Therefore, the rate of cytosolic MCC formation is calculated as:

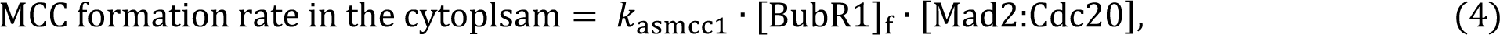

where,

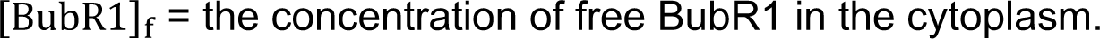

#### Stage 2: Effect of MCC formation on the timing of metaphase-to-anaphase transition

We used a previously described model of metaphase-to-anaphase transition to simulate the effect of the MCC generated on the duration of mitosis (Chen et al., 2019; He et al., 2011). In this model (schematic at the top of Figure S1B), Cyclin B (‘CycB’) is synthesized at a constant rate and degraded upon APC:Cdc20-dependent ubiquitination (denoted simply as Cdc20). The abundance of CycB determines the activity of CDK1:CycB complexes, which in turn determines the activity of the eSAC complexes via phosphorylation. CDK1:CycB activity is antagonized by the counter-acting protein phosphatase PP2A:B56 (‘CAPP’) (Bouchoux and Uhlmann, 2011; Sullivan et al., 2004). This scheme is consistent with recent data revealing that CDK1:CycB phosphorylates Bub1 to promote its interaction with Mad1 (Ji et al., 2017). Furthermore, Mps1 kinase activity is also downregulated by PP2A (Espert et al., 2014; Hayward et al., 2019). This scheme regulates the amount of active eSAC.

The active eSAC ultimately produces MCC according to the scheme discussed in detail above with BubR1 as an MCC component. Therefore, we modified the original model to include BubR1 as well as the dissociation of MCC into its constituent proteins (shown by the red dashed arrow in Figure S1B). Furthermore, active APC:Cdc20 promotes the inactivation of closed/active Mad2 in MCC; this positive feedback of active Cdc20 on its own release from the MCC accelerates the activation of APC:Cdc20 during the transition into anaphase (Chen et al., 2019; He et al., 2011).

The equations for this model are given below using the following notation:

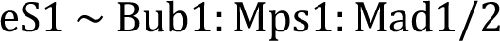

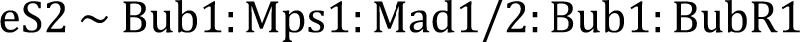

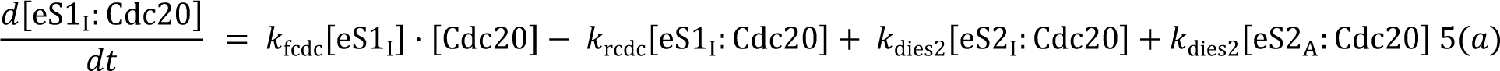

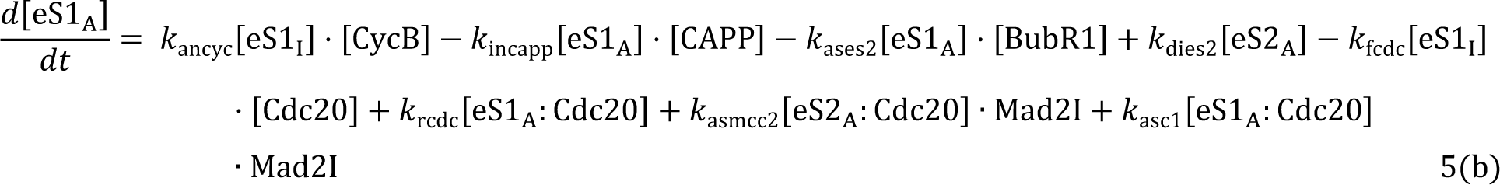

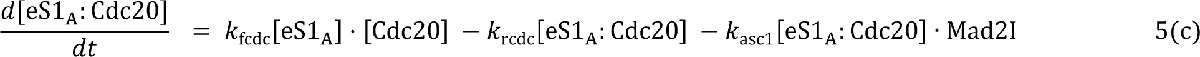

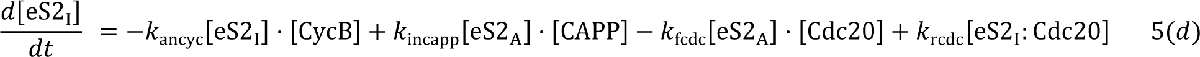

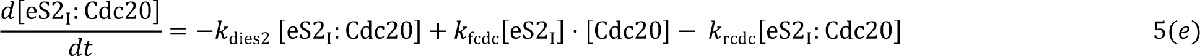

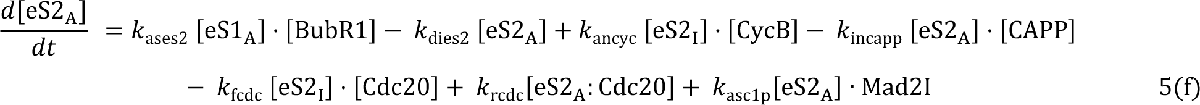

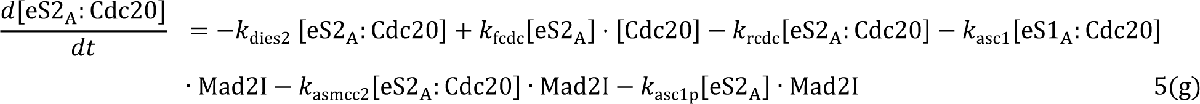

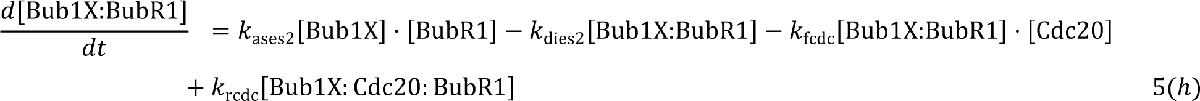

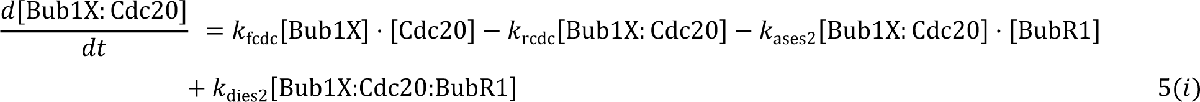

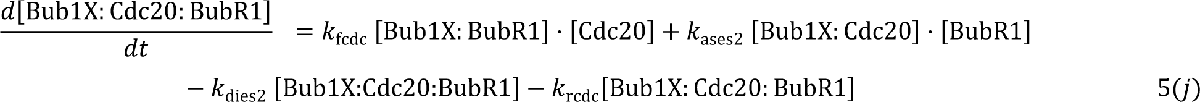

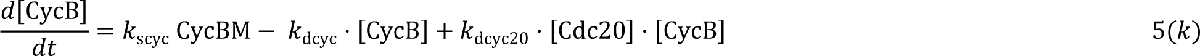

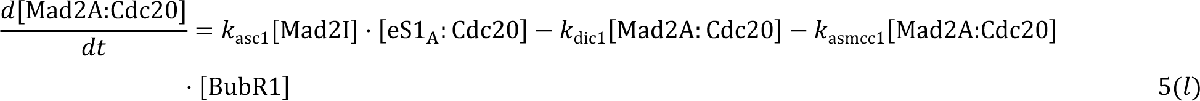

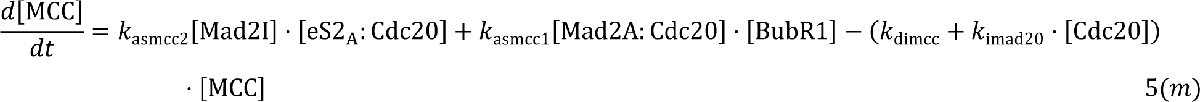

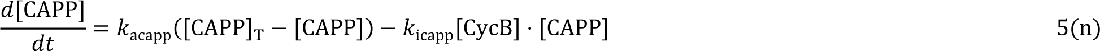

The molecular species, chemical reactions, parameters, and constraints involved in the model are presented in an Excel file and Table S1 (Supporting Information). A MATLAB script was used to read this file and produce a file with Ordinary Differential Equations (ODEs) describing the rate of change of all molecular species. We numerically integrated the ODEs to calculate the time evolution of [CycB] and the other molecular species. For the initial conditions of species involving BubR1 and Cdc20 bound to Bub1 phosphodomain (both signaling and non-signaling) we used the equilibrium concentrations. These concentrations depend on the concentration of eSAC activator complex and were calculated at the start of each simulation. The initial concentrations of the following species were kept constant: [CycB] = 45 nM, [Mad2:Cdc20] = 0 nM, [MCC] = 25 nM, [CAPP] = 5 nM. In the simulation, the system evolves towards a steady state corresponding to anaphase (low [CycB] and high [Cdc20]). We assume that a cell exits mitosis when [CycB] drops below 1 nM (Fig. S1E). If the initial conditions are metaphase-like (high [CycB] and low [Cdc20]), small variations in the initial conditions does not qualitatively affect the outcome of the model. Furthermore, the main result of this analysis – Bub1-BubR1 produces MCC at a higher rate than Bub1 – is robust, even though many different combinations of parameters produce similar looking dose:response curves.

#### Modeling the activities of eSAC phosphodomains containing four MELT motifs and the KI motifs

This model simulates the dose-response data for the eSAC systems involving MELT and KI motifs. As before, the simulation takes place in two stages. In the first stage, we calculate the steady-state concentrations of SAC signaling proteins (Bub1 and BubR1) bound to the phosphorylated MELT motifs and KI motifs of an eSAC phosphodomain. This is followed by calculation of MCC generation using the steady-state concentrations of MELpT:Bub1 and MELpT:Bub1:BubR1. In the second stage, we simulate the duration of mitosis, according to the overall reaction scheme developed by He et al.

#### Stage 1: Simulation of SAC protein recruitment by the eSAC phosphodomains

##### Rules for protein-protein interactions

The phosphodomain consists of four MELT motifs and either active or inactive KI motifs. All four MELT motifs in phosphodomains complexed with the Mps1 kinase domain are assumed to be phosphorylated. They can recruit Bub1, which represents Bub1-Bub3 in this model. Upon binding to the MELT motif, Bub1 recruits BubR1 representing BubR1-Bub3. The KI1 motif can bind to Bub1, whereas the KI2 motif can bind BubR1. We assume that MELT and KI motifs in the eSAC phosphodomain interact with SAC proteins independently. The KI-bound Bub1 and BubR1 do not participate in SAC signaling. Therefore, the KI motifs act as sinks that reduce free Bub1 and BubR1 concentrations. We assigned the same rate of binding of the ‘Bub1’ protein to each MELT motif (*kf* in Table S2), but assigned a low unbinding rate (*kr*, Table S2) for the strong MELT motifs (MELT 1, 12 and 14), and a higher unbinding rate for the weak MELT motif (MELT 13) following previous studies (Chen et al., 2019; Vleugel et al., 2015). We also chose the dissociation constant for KI1:Bub1 binding to be equal to the dissociation constant for the KI2:BubR1 binding.

##### Calculation of the steady-state concentrations of signaling complexes

This model avoids unnecessary complexity by assuming that the rate of MCC formation is simply proportional to the number of phosphorylated MELT motifs that recruit Bub1 or both Bub1 and BubR1. This simplification is justified, because the recruitment of Mad1, Mad2, and Cdc20 is independent of BubR1 recruitment. Thus, the main goal is to determine the steady-state concentrations of the two distinct signaling complexes: MELpT:Bub1 or MELpT:Bub1:BubR1. This calculation is performed as follows.

Each phosphorylated MELT motif can be in one of three possible states: MELpT (unbound MELT), MELpT:Bub1 (MELT bound by Bub1), and MELpT:Bub1:BubR1 (MELT bound by Bub1 and BubR1). Since there are four MELT motifs in each eSAC activator complex, the number of possible states for the phosphomimic become 3^4^ = 81. The time evolution of concentrations of different Bub1 and BubR1 bound states of eSAC activator complex is given by:

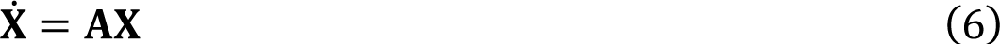

where ***X** = {x_1_, x_2_, …, x_N_}* is a vector of concentrations of the *N* = 81 different Bub1 and BubR1 binding states of the phosphodomain, and **A** is the rate matrix.

Similarly, each KI motif of the phosphodomain can be in two states: bound or unbound. The binding of Bub1 and BubR1 to KI motifs is described by the set of equations:

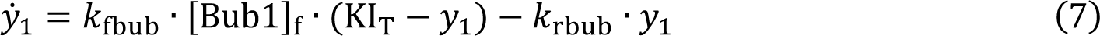

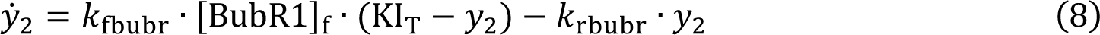

where, KI_T_ is the total concentration of KI motifs, and

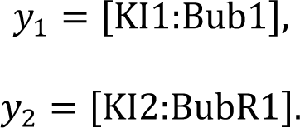

[Bub1]_f_ and [BubR1]_f_ are the cytoplasmic concentrations of free Bub1 and BubR1, respectively. The parameters k_fbub_ and k_fbub_ are the binding and unbinding rate constants between Bub1 and the first KI1, and the parameters k_fbubr_ and k_fbubr_ are the binding and unbinding rates between BubR1 and KI2.

The concentrations satisfy the constraints:

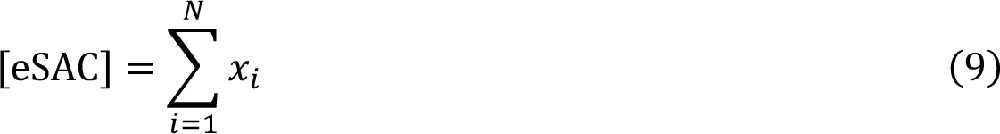

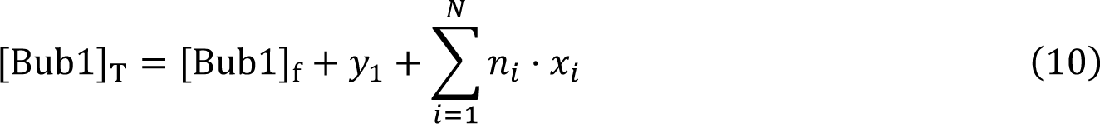

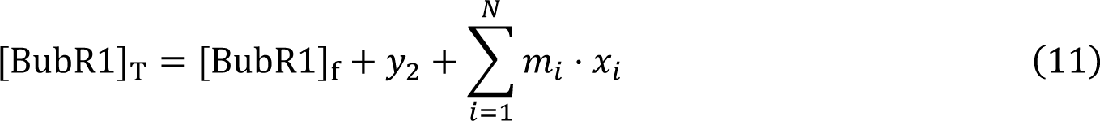

Here, *x_i_* is the concentration of the i-th species, *n_i_* = number of Bub1 bound to i-th species of the phosphodomain, and *m_i_* = number of BubR1 units bound to the i-th species. We assume [Bub1]_T_ = 100 nM and [BubR1]_T_ = 100 nM. The equilibrium concentration of each state was obtained by numerically solving

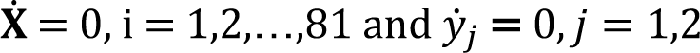

In experiments, the concentration of eSAC activator complex is measured in arbitrary units of mCherry fluorescence (a. u.), whereas in our model the unit of concentration is nano moles (nM). In our simulations, we chose the maximum value of concentration of eSAC activator complex (the value corresponding to 20 a. u. in experiments) to be 200 nM. For easier comparison to experimental figures, in our simulation results the eSAC activator complex concentration is expressed in arbitrary units, with 1 a. u. of fluorescence corresponding to 10 nM.

Figure S4A shows the abundance of different Bub1 and BubR1-bound states as functions of the total concentration of eSAC activator complex for KI1*-KI2*. At low eSAC concentrations, the eSAC tends to be highly loaded, with Bub1 and BubR1 on every MELT motif. However, for cells with a high eSAC concentration, [eSAC] >> [Bub1]_T_, the most abundant eSAC species is one that does not bind any Bub1 at all (not shown), followed by species that bind either Bub1 or Bub1-BubR1 at only one of the four MELT motifs. We define the sum of concentration of MELpT:Bub1 and MELpT:Bub1:BubR1 as [eSAC]_T_:

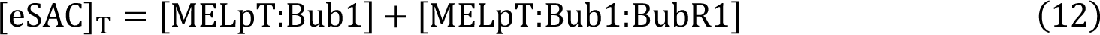

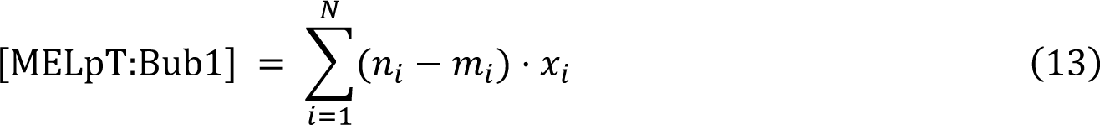

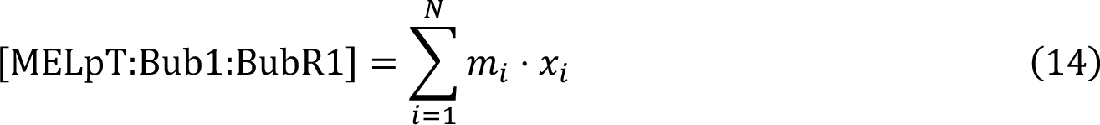

### Formation of MCC by the eSAC signaling complexes

We assume that the MELpT:Bub1 and MELpT:Bub1:BubR1 complexes catalyze the assembly of MCC at the apparent rates constants *k_MCC_* and *k’_MCC_*. A schematic diagram of the molecular mechanism underlying this model is displayed in Figure 4A.

We assume that the recruitment of SAC proteins (Bub1 and BubR1) to MELT motifs of the eSAC activator complexes enables the incorporation of Cd20 into MCC. Since different species catalyze this reaction at different rates, we define the effective rate of conversion, *k_asmcc_*, as the concentration-weighted sum of the conversion rates of each eSAC complex:

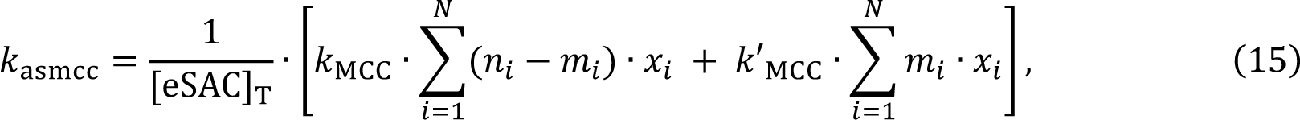

where *k_MCC_* and *k’_MCC_* are the MCC formation rates due to MELpT:Bub1 and MELpT:Bub1:BubR1, respectively. Note, for most species, the rate of MCC generation is additive. For example, *k_0012_ = k_MCC_ + k’_MCC_* for a phosphodomain that binds only Bub1 at the 13^th^ MELT motif and Bub1:BubR1 at 14^th^ MELT motif (Table S3). Using [eSAC]_T_ and k_asmcc_ as inputs for the He model (discussed below), we calculated the time evolution of Cyclin B concentration and from it the time in mitosis.

#### Stage 2: The effect of MCC produced on exit from mitosis

To calculate the effect of MCC generated by the eSAC on mitotic exit, we used a simplified version of the model of the mitotic checkpoint proposed by the He model (He et al., 2011). Active eSAC signaling complexes (eSAC_A_) generate MCC, as described in the previous section. The temporal dynamics of our mitotic checkpoint model are determined by the ordinary differential equations.

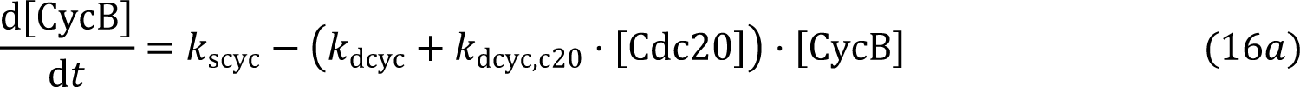

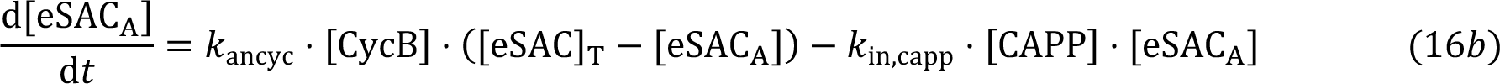

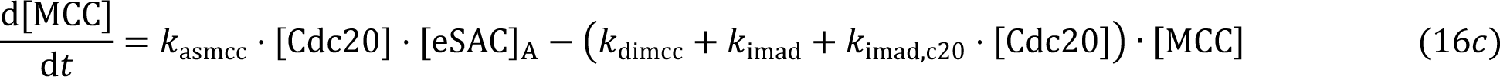

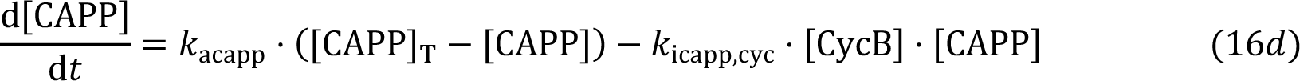

In these equations, [CycB] = [CDK:CycB], [Cdc20] = [APC:Cdc20], [eSAC]_T_ is the total concentration of Bub1 and BubR1-bound eSAC signaling complexes (Eq. 10), which is either in the active, signaling-competent state eSAC_A_, or the inactive state, [eSAC_I_] = [eSAC]_T_ − [eSACA]. [MCC] and [CAPP] refer to the concentrations of the mitotic checkpoint complex and the CDK-counteracting protein phosphatase, respectively. In addition, the total concentration of Cdc20 is:

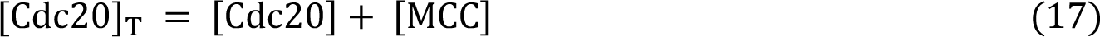

The values of the parameters in the model and of the fixed concentrations of some components, as listed in Table S2, are taken from (He et al., 2011).

### Simulation of time in mitosis

To determine the timing of the metaphase-to-anaphase transition, we assume that a cell exits mitosis when [CycB] drops below 1 nM. We numerically integrated Ordinary Differential Equations (ODEs) to calculate the time evolution of [CycB] and the components of the cell cycle machinery. As before, the initial conditions for the simulation are chosen to be [CycB] = 45 nM, [eSAC_A_] = 0, [MCC] = 25 nM, and [CAPP] = 5 nM. The qualitative aspects of our results do not depend on the initial conditions. Figure S3C displays typical time courses for [CycB], for different eSACs, for [eSAC activator complex] = 10 a. u. The system always comes out of mitosis (as seen by the drop in [CycB]), albeit after different time delays.

---

**Figure S1.**
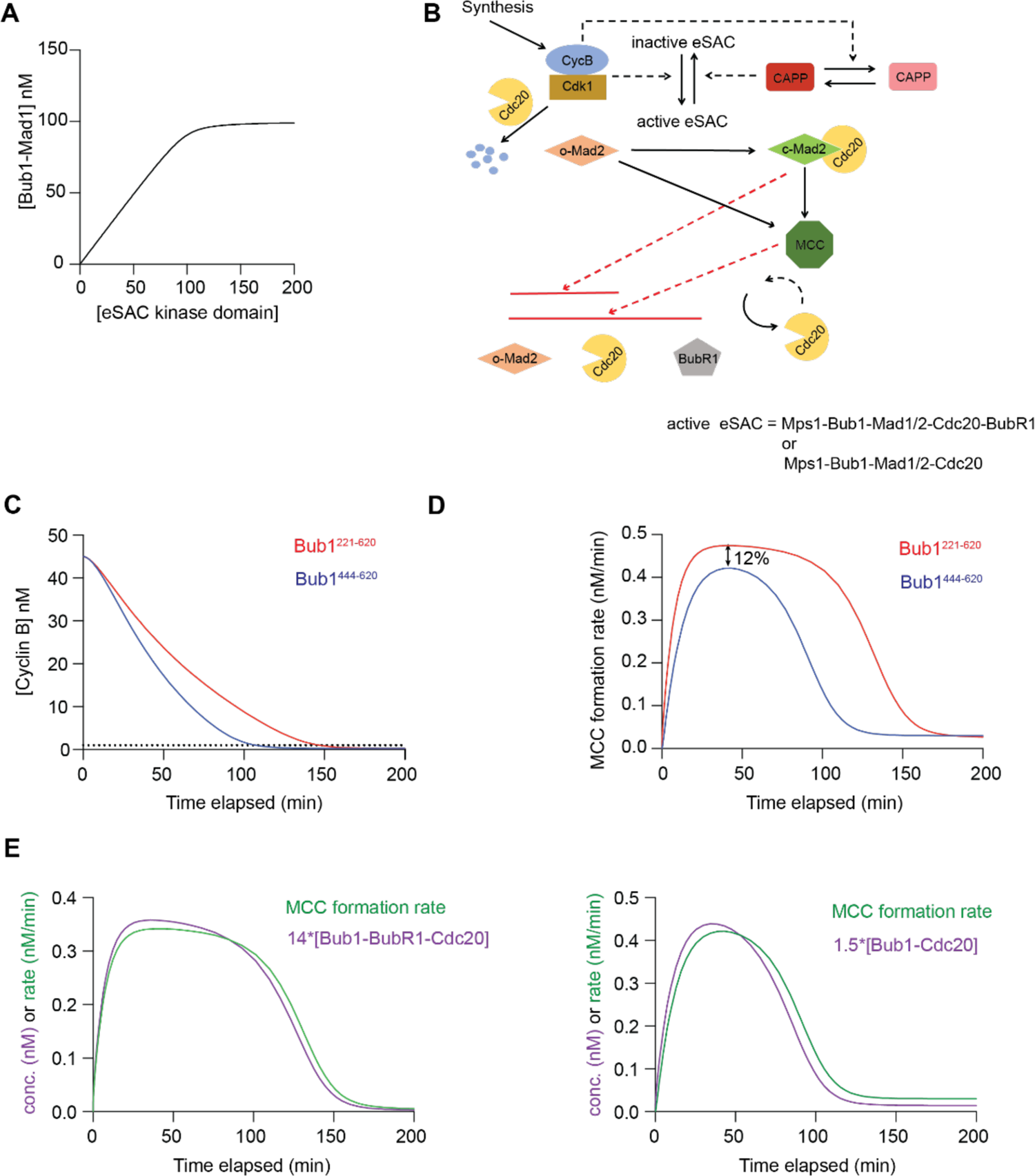
Simulation of the dose-response curves for the Bub1 eSAC system. **A** The amount of the eSAC signaling complex (Bub1-Mad1) saturates due to the limited availability of Mad1 (100 nM). **B** Schematic of the model used to simulate the dose-response data for the “full” and “truncated” Bub1 phosphodomain used in the eSAC system. **C** Temporal evolution of Cyclin B concentration for the two eSAC systems. **D** Assessment of MCC formation rates for the full and truncated phosphodomain. **E** Superimposition of the dynamics of the MCC formation rate and the concentration of signaling complexes that produce MCC. Note that the ∼ 10-fold difference in the scaling in the two panels used for displaying with clarity the concentrations of the signaling complex.

**Supplementary Figure 2.**
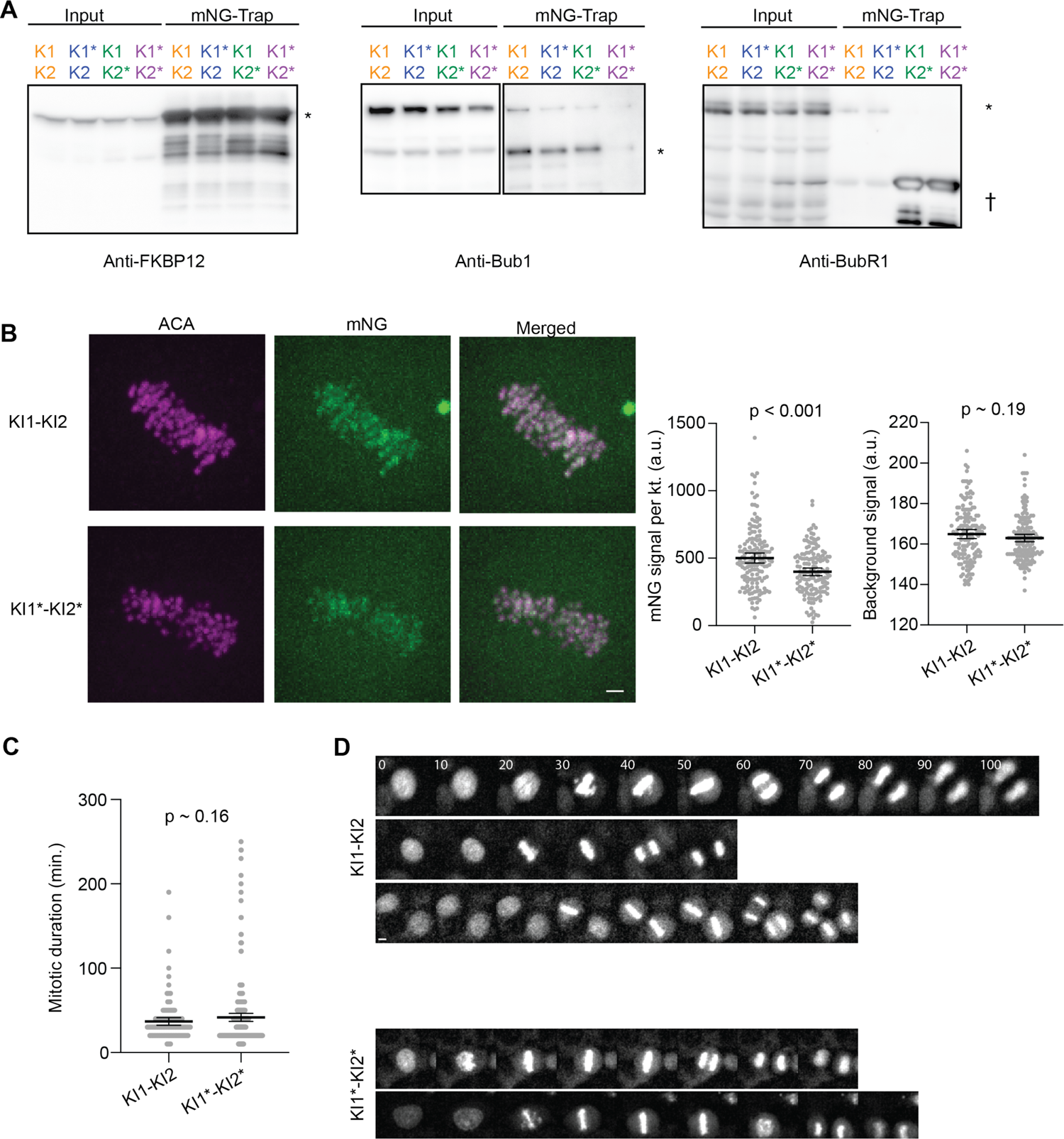
Characterization of the interactions of the KI-motif containing phosphodomains with Bub1 and BubR1 and their cellular localization. **A** Immunoblots probing for the indicated protein for assessing the co-immunoprecipitation of Bub1 and BubR1 with the indicated eSAC phosphodomain (at the top). The asterisk marks the position of the band corresponding to the full-length protein. The cross marks bands may be degradation products of BubR1. **B** Left: Representative micrographs of metaphase cells displaying the indicated proteins (scale bar ∼ 1.1 μm). Right: Quantification of kinetochore-localized mNG signal corresponding to the indicated phosphodomain (n = 149, 147 for KI1-KI2 and KI1*-KI2* respectively). The scatterplot on the right displays the background intensity measured in a 12×12 pixel box concentric with the 6×6 box corresponding to the kinetochore spot. The p-values indicate the results of Welch’s t-test. Similar mean values of the background fluorescence, which is indicative of the expression level of the phosphodomain, indicate that the difference in the kinetochore-localized phosphodomain is not due to a difference in the expression level. **C** Duration of mitosis of untreated cells expressing KI1-KI2-mNG-2xFkbp12 or KI1*-KI2*-mNG-2xFkbp12 along with Frb-mCherry-Mps1^500-817^ in the absence of Rapamycin (n = 124, 175 from 2 experiments for KI1-KI2 and KI1*-KI2* respectively). The p-value indicates the result of Welch’s t-test. **D** Selected montages show mitotic progression of the indicated HeLa cell lines transiently transfected with mNG-H2B expression plasmids. (scale bar ∼ 4.87 μm). Numbers at the top left corner indicate the time elapsed in minutes from the first frame.

**Figure S3.**
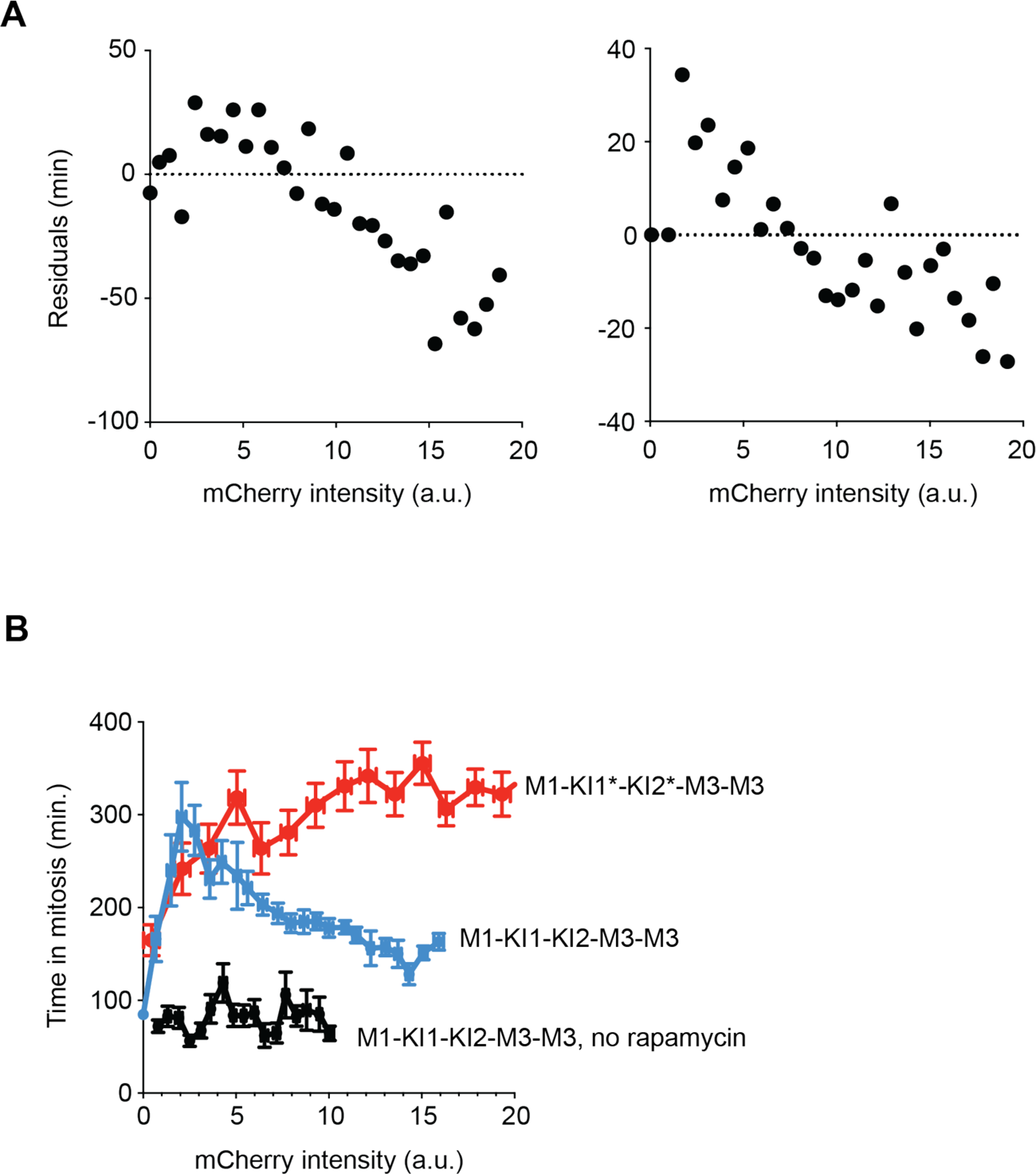
Non-monotonic nature of the dose-response curves and dose-response data for phosphodomains containing the two KI motifs along with seven MELT motifs. **A** Residual plots from a 4-parameter sigmoidal fit to the dose-response data for the KI1*-KI2* (left) and KI1-KI2* (right) phosphodomains. **B** Dose-response data for the indicated phosphodomains. Circles represent average values of binned data (vertical and horizontal lines display standard error on each mean value. n= 498, 1313, and 394 respectively for M1-KI1-KI2-M3-M3, M1-KI1*-KI2*-M3-M3, and M1-KI1-KI2-M3-M3 without Rapamycin respectively pooled from two or more experiments).

**Supplementary Figure S4.**
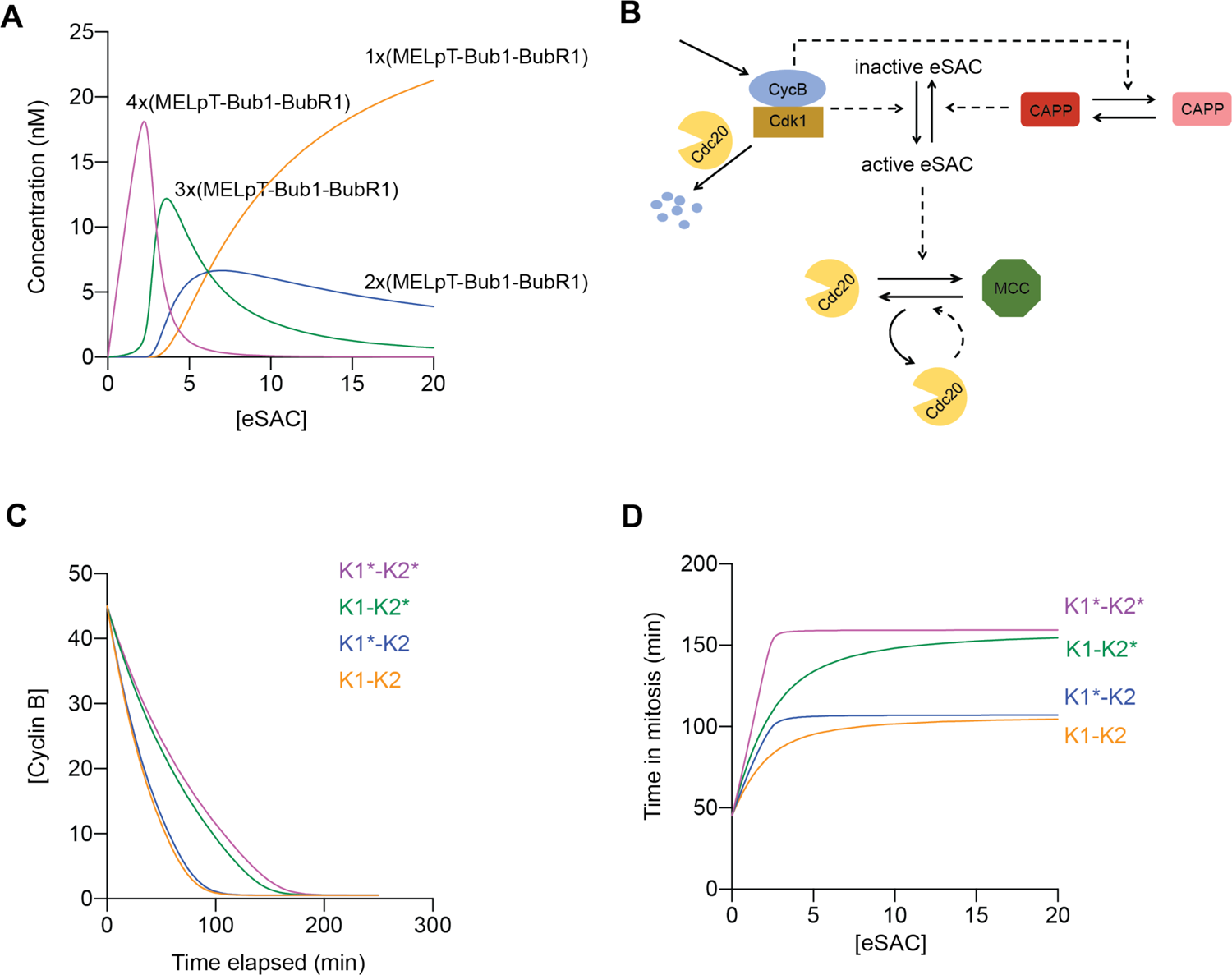
Simulation of the dose-response data for eSAC systems containing the KI motifs and MELT motifs. **A** Abundance of the indicated eSAC signaling complex (of the form n*(MELpT-Bub1-BubR1)) as a function of the total concentration of the eSAC activator complex. The number n indicates the number of MELT motifs bound to Bub1-BubR1. Also see Table S3. **B** Schematic of the model used to simulate the dose-response data for eSAC with MELT and KI motifs. **C** Temporal evolution of Cyclin B concentration for the MELT and KI motif eSAC system. **D** Simulation of dose-response curves for the four phosphodomains without synergistic interactions between the MELT motifs.

## References

1. Aravamudhan, P., A.A. Goldfarb, and A.P. Joglekar. 2015. The kinetochore encodes a mechanical switch to disrupt spindle assembly checkpoint signalling. Nat Cell Biol. 17:868–879.

2. Bolanos-Garcia, V.M., T. Lischetti, D. Matak-Vinkovic, E. Cota, P.J. Simpson, D.Y. Chirgadze, D.R. Spring, C.V. Robinson, J. Nilsson, and T.L. Blundell. 2011a. Structure of a Blinkin-BUBR1 complex reveals an interaction crucial for kinetochore-mitotic checkpoint regulation via an unanticipated binding Site. Structure. 19:1691–1700.

3. Bolanos-Garcia, Victor M., T. Lischetti, D. Matak-Vinković, E. Cota, Pete J. Simpson, Dimitri Y. Chirgadze, David R. Spring, Carol V. Robinson, J. Nilsson, and Tom L. Blundell. 2011b. Structure of a Blinkin-BUBR1 Complex Reveals an Interaction Crucial for Kinetochore-Mitotic Checkpoint Regulation via an Unanticipated Binding Site. Structure. 19:1691–1700.

4. Bouchoux, C., and F. Uhlmann. 2011. A quantitative model for ordered Cdk substrate dephosphorylation during mitotic exit. Cell. 147:803–814.

5. Chen, C., I.P. Whitney, A. Banerjee, C. Sacristan, P. Sekhri, D.M. Kern, A. Fontan, G. Kops, J.J. Tyson, I.M. Cheeseman, and A.P. Joglekar. 2019. Ectopic Activation of the Spindle Assembly Checkpoint Signaling Cascade Reveals Its Biochemical Design. Curr Biol. 29:104–119 e110.

6. Collin, P., O. Nashchekina, R. Walker, and J. Pines. 2013. The spindle assembly checkpoint works like a rheostat rather than a toggle switch. Nat Cell Biol. 15:1378–1385.

7. Di Fiore, B., Norman E. Davey, A. Hagting, D. Izawa, J. Mansfeld, Toby J. Gibson, and J. Pines. 2015. The ABBA Motif Binds APC/C Activators and Is Shared by APC/C Substrates and Regulators. Developmental Cell. 32:358–372.

8. Dick, A.E., and D.W. Gerlich. 2013. Kinetic framework of spindle assembly checkpoint signalling. Nat Cell Biol. 15:1370–1377.

9. Elowe, S., S. Hummer, A. Uldschmid, X. Li, and E.A. Nigg. 2007. Tension-sensitive Plk1 phosphorylation on BubR1 regulates the stability of kinetochore microtubule interactions. Genes Dev. 21:2205–2219.

10. Espert, A., P. Uluocak, R.N. Bastos, D. Mangat, P. Graab, and U. Gruneberg. 2014. PP2A-B56 opposes Mps1 phosphorylation of Knl1 and thereby promotes spindle assembly checkpoint silencing. J Cell Biol. 206:833–842.

11. Faesen, A.C., M. Thanasoula, S. Maffini, C. Breit, F. Muller, S. van Gerwen, T. Bange, and A. Musacchio. 2017. Basis of catalytic assembly of the mitotic checkpoint complex. Nature.

12. Foley, E.A., M. Maldonado, and T.M. Kapoor. 2011. Formation of stable attachments between kinetochores and microtubules depends on the B56-PP2A phosphatase. Nat Cell Biol. 13:1265–1271.

13. Hayward, D., J. Bancroft, D. Mangat, T. Alfonso-Perez, S. Dugdale, J. McCarthy, F.A. Barr, and U. Gruneberg. 2019. Checkpoint signaling and error correction require regulation of the MPS1 T-loop by PP2A-B56. J Cell Biol. 218:3188–3199.

14. He, E., O. Kapuy, R.A. Oliveira, F. Uhlmann, J.J. Tyson, and B. Novák. 2011. System-level feedbacks make the anaphase switch irreversible. Proceedings of the National Academy of Sciences.

15. Ji, Z., H. Gao, L. Jia, B. Li, and H. Yu. 2017. A sequential multi-target Mps1 phosphorylation cascade promotes spindle checkpoint signaling. Elife. 6.

16. Jia, L., B. Li, and H. Yu. 2016. The Bub1-Plk1 kinase complex promotes spindle checkpoint signalling through Cdc20 phosphorylation. Nat Commun. 7:10818.

17. Kern, D.M., T. Kim, M. Rigney, N. Hattersley, A. Desai, and I.M. Cheeseman. 2015. The outer kinetochore protein KNL-1 contains a defined oligomerization domain in nematodes. Mol Biol Cell. 26:229–237.

18. Khandelia, P., K. Yap, and E.V. Makeyev. 2011. Streamlined platform for short hairpin RNA interference and transgenesis in cultured mammalian cells. Proc Natl Acad Sci USA. 108:12799–12804.

19. Kiyomitsu, T., H. Murakami, and M. Yanagida. 2011. Protein interaction domain mapping of human kinetochore protein Blinkin reveals a consensus motif for binding of spindle assembly checkpoint proteins Bub1 and BubR1. Mol Cell Biol. 31:998–1011.

20. Krenn, V., K. Overlack, I. Primorac, S. van Gerwen, and A. Musacchio. 2014. KI motifs of human Knl1 enhance assembly of comprehensive spindle checkpoint complexes around MELT repeats. Curr Biol. 24:29–39.

21. Krenn, V., A. Wehenkel, X. Li, S. Santaguida, and A. Musacchio. 2012. Structural analysis reveals features of the spindle checkpoint kinase Bub1-kinetochore subunit Knl1 interaction. J Cell Biol. 196:451–467.

22. Kruse, T., G. Zhang, M.S. Larsen, T. Lischetti, W. Streicher, T. Kragh Nielsen, S.P. Bjorn, and J. Nilsson. 2013. Direct binding between BubR1 and B56-PP2A phosphatase complexes regulate mitotic progression. J Cell Sci. 126:1086–1092.

23. Lara-Gonzalez, P., T. Kim, K. Oegema, K. Corbett, and A. Desai. 2021a. A tripartite mechanism catalyzes Mad2-Cdc20 assembly at unattached kinetochores. Science. 371:64–67.

24. Lara-Gonzalez, P., J. Pines, and A. Desai. 2021b. Spindle assembly checkpoint activation and silencing at kinetochores. Semin. Cell Dev. Biol. 117:86–98.

25. Leontiou, I., N. London, K.M. May, Y. Ma, L. Grzesiak, B. Medina-Pritchard, P. Amin, A.A. Jeyaprakash, S. Biggins, and K.G. Hardwick. 2019. The Bub1-TPR Domain Interacts Directly with Mad3 to Generate Robust Spindle Checkpoint Arrest. Curr Biol. 29:2407–2414 e2407.

26. Musacchio, A. 2015. The Molecular Biology of Spindle Assembly Checkpoint Signaling Dynamics. Curr Biol. 25:R1002–1018.

27. Nijenhuis, W., G. Vallardi, A. Teixeira, G.J. Kops, and A.T. Saurin. 2014. Negative feedback at kinetochores underlies a responsive spindle checkpoint signal. Nat Cell Biol. 16:1257–1264.

28. Overlack, K., T. Bange, F. Weissmann, A.C. Faesen, S. Maffini, I. Primorac, F. Muller, J.M. Peters, and A. Musacchio. 2017. BubR1 Promotes Bub3-Dependent APC/C Inhibition during Spindle Assembly Checkpoint Signaling. Curr Biol. 27:2915–2927 e2917.

29. Overlack, K., I. Primorac, M. Vleugel, V. Krenn, S. Maffini, I. Hoffmann, G.J. Kops, and A. Musacchio. 2015. A molecular basis for the differential roles of Bub1 and BubR1 in the spindle assembly checkpoint. Elife. 4:e05269.

30. Piano, V., A. Alex, P. Stege, S. Maffini, G.A. Stoppiello, P.J. Huis In’t Veld, I.R. Vetter, and A. Musacchio. 2021. CDC20 assists its catalytic incorporation in the mitotic checkpoint complex. Science. 371:67-71.

31. Primorac, I., J.R. Weir, E. Chiroli, F. Gross, I. Hoffmann, S. van Gerwen, A. Ciliberto, and A. Musacchio. 2013. Bub3 reads phosphorylated MELT repeats to promote spindle assembly checkpoint signaling. Elife. 2:e01030.

32. Qian, J., M.A. Garcia-Gimeno, M. Beullens, M.G. Manzione, G. Van der Hoeven, J.C. Igual, M. Heredia, P. Sanz, L. Gelens, and M. Bollen. 2017. An Attachment-Independent Biochemical Timer of the Spindle Assembly Checkpoint. Mol Cell. 68:715–730 e715.

33. Roy, B., S.J.Y. Han, A.N. Fontan, S. Jema, and A.P. Joglekar. 2022. Aurora B phosphorylates Bub1 to promote spindle assembly checkpoint signaling. Curr Biol. 32:237–247 e236.

34. Roy, B., S.J.Y. Han, A.N. Fontan, and A.P. Joglekar. 2020. The copy-number and varied strength of MELT motifs in Spc105 balance the strength and responsiveness of the Spindle Assembly Checkpoint. Elife. 9.

35. Saurin, A.T., M.S. van der Waal, R.H. Medema, S.M. Lens, and G.J. Kops. 2011. Aurora B potentiates Mps1 activation to ensure rapid checkpoint establishment at the onset of mitosis. Nat Commun. 2:316.

36. Suijkerbuijk, S.J., M. Vleugel, A. Teixeira, and G.J. Kops. 2012. Integration of kinase and phosphatase activities by BUBR1 ensures formation of stable kinetochore-microtubule attachments. Dev Cell. 23:745–755.

37. Sullivan, M., T. Higuchi, V.L. Katis, and F. Uhlmann. 2004. Cdc14 phosphatase induces rDNA condensation and resolves cohesin-independent cohesion during budding yeast anaphase. Cell. 117:471–482.

38. Taylor, S.S., E. Ha, and F. McKeon. 1998. The human homologue of Bub3 is required for kinetochore localization of Bub1 and a Mad3/Bub1-related protein kinase. J Cell Biol. 142:1–11.

39. Taylor, S.S., D. Hussein, Y. Wang, S. Elderkin, and C.J. Morrow. 2001. Kinetochore localisation and phosphorylation of the mitotic checkpoint components Bub1 and BubR1 are differentially regulated by spindle events in human cells. J Cell Sci. 114:4385–4395.

40. Tromer, E., D. Bade, B. Snel, and G.J. Kops. 2016. Phylogenomics-guided discovery of a novel conserved cassette of short linear motifs in BubR1 essential for the spindle checkpoint. Open Biol. 6.

41. Vleugel, M., M. Omerzu, V. Groenewold, M.A. Hadders, S.M. Lens, and G.J. Kops. 2015. Sequential multisite phospho-regulation of KNL1-BUB3 interfaces at mitotic kinetochores. Mol Cell. 57:824–835.

42. Vleugel, M., E. Tromer, M. Omerzu, V. Groenewold, W. Nijenhuis, B. Snel, and G.J. Kops. 2013. Arrayed BUB recruitment modules in the kinetochore scaffold KNL1 promote accurate chromosome segregation. J Cell Biol. 203:943–955.

43. von Schubert, C., F. Cubizolles, Jasmine M. Bracher, T. Sliedrecht, Geert J.P.L. Kops, and Erich A. Nigg. 2015. Plk1 and Mps1 Cooperatively Regulate the Spindle Assembly Checkpoint in Human Cells. Cell Reports. 12:66–78.

44. Wang, J., Z. Wang, T. Yu, H. Yang, D.M. Virshup, G.J.P.L. Kops, S.H. Lee, W. Zhou, X. Li, W. Xu, and Z. Rao. 2016a. Crystal structure of a PP2A B56-BubR1 complex and its implications for PP2A substrate recruitment and localization. Protein & Cell. 7:516–526.

45. Wang, X., R. Bajaj, M. Bollen, W. Peti, and R. Page. 2016b. Expanding the PP2A Interactome by Defining a B56-Specific SLiM. Structure. 24:2174–2181.

46. Yuan, I., I. Leontiou, P. Amin, K.M. May, S. Soper Ni Chafraidh, E. Zlamalova, and K.G. Hardwick. 2017. Generation of a Spindle Checkpoint Arrest from Synthetic Signaling Assemblies. Curr Biol. 27:137–143.

47. Zhang, G., T. Lischetti, D.G. Hayward, and J. Nilsson. 2015. Distinct domains in Bub1 localize RZZ and BubR1 to kinetochores to regulate the checkpoint. Nat Commun. 6:7162.

48. Zhang, G., T. Lischetti, and J. Nilsson. 2014. A minimal number of MELT repeats supports all the functions of KNL1 in chromosome segregation. J Cell Sci. 127:871–884.

49. Zhang, G., B.L. Mendez, G.G. Sedgwick, and J. Nilsson. 2016. Two functionally distinct kinetochore pools of BubR1 ensure accurate chromosome segregation. Nat Commun. 7:12256.

