## Supplementary Tables for "BubR1 recruitment to the kinetochore via Bub1 enhances Spindle Assembly Checkpoint signaling"

**Table S1: Parameters used for simulating the eSAC response consisting of Bub1-BubR1 heterodimer.**

| Parameter | Description | Value |
|---|---|---|
| $k_{fbubr}$ | Rate of BubR1 binding to Bub1 | 0.005 $nM^{-1} \cdot min^{-1}$ |
| $k_{rbubr}$ | Rate of BubR1 dissociation from Bub1 | 1 $min^{-1}$ |
| $k_{scyc}$ | Rate of Cyclin B synthesis | 0.05 $nM \cdot min^{-1}$ |
| $k_{dcyc}$ | Rate of Cyclin B degradation | 0.001 $min^{-1}$ |
| $k_{dcyc20}$ | Rate of APC:Cdc20 mediated degradation of Cyclin B | 0.04 $min^{-1}$ |
| $k_{ancyc}$ | Rate of Cdk1:CycB catalyzed activation of inactive eSAC | 1 $nM^{-1} \cdot min^{-1}$ |
| $k_{incapp}$ | Rate of CAPP catalyzed inactivation of active eSAC | 1 $nM^{-1} \cdot min^{-1}$ |
| $k_{asc1}$ | Rate of Mad2:Cdc20 formation | 0.5 $nM^{-1} \cdot min^{-1}$ |
| $k_{idic1}$ | Rate of dissociation of Mad2:Cdc20 | 1 $min^{-1}$ |
| $k_{imad20}$ | Rate of Cdc20 mediated inactivation of MCC complex | 0.008 $nM^{-1} \cdot min^{-1}$ |
| $k_{asmcc1}$ | Rate of Mad2:Cdc20 and BubR1 binding in cytosol | 0.002 $nM^{-1} \cdot min^{-1}$ |
| $k_{asmcc2}$ | Rate of MCC formation at the phosphodomain | 0.5 $nM^{-1} \cdot min^{-1}$ |
| $k_{dimcc}$ | Rate of dissociation of MCC complex | 0.02 $min^{-1}$ |
| $k_{acapp}$ | Rate of CAPP dephosphorylation | 0.1 $min^{-1}$ |
| $k_{icapp}$ | Rate of Cdk1 catalyzed phosphorylation of CAPP | 0.02 $nM^{-1} \cdot min^{-1}$ |

| | | |
|---|---|---|
| $k_{fcdc}$ | Rate of Cdc20 binding to Bub1 | 0.12 $nM^{-1}\cdot min^{-1}$ |
| $k_{rcdc}$ | Rate of dissociation of Bub1:Cdc20 complex | 5 $min^{-1}$ |
| $[Bub1]_T$ | Total concentration of Bub1 phosphodomain | 200 nM |
| $[BubR1]_T$ | Total concentration of BubR1 | 80 nM |
| $[CycB]_M$ | Concentration of Cyclin B in mitosis | 50 nM |
| $[Cdc20]_T$ | Total concentration of Cdc20 | 25 nM |
| $[Mad2]_T$ | Total concentration of Mad2 | 50 nM |
| $[CAPP]_T$ | Total concentration of CAPP (a Cdk1 counteracting protein phosphatase) | 50 nM |

**Table S2 Parameters used for simulating eSAC activity and response of the SAC machinery**

| Parameter | Description | Value |
|---|---|---|
| $k_{f1}$ | Rate of Bub1 binding to MELT motif 1 | 1 $nM^{-1}$ $min^{-1}$ |
| $k_{f12}$ | Rate of Bub1 binding to MELT motif 12 | 1 $nM^{-1}$ $min^{-1}$ |
| $k_{f13}$ | Rate of Bub1 binding to MELT motif 13 | 1 $nM^{-1}$ $min^{-1}$ |
| $k_{f14}$ | Rate of Bub1 binding to MELT motif 14 | 1 $nM^{-1}$ $min^{-1}$ |
| $k_{r1}$ | Rate of Bub1 dissociation from MELT motif 1 | 0.1 $min^{-1}$ |
| $k_{r12}$ | Rate of Bub1 dissociation from MELT motif 12 | 0.1 $min^{-1}$ |
| $k_{r13}$ | Rate of Bub1 dissociation from MELT motif 13 | 1 $min^{-1}$ |
| $k_{r14}$ | Rate of Bub1 dissociation from MELT motif 14 | 0.1 $min^{-1}$ |
| $k_{fbubr}$ | Rate of BubR1 binding to Bub1 | 1 $nM^{-1}$ $min^{-1}$ |
| $k_{rbubr}$ | Rate of BubR1 dissociation from Bub1 | 0.1 $min^{-1}$ |
| $ki_{f1}$ | Rate of Bub1 binding to KI1 motif | 1 $nM^{-1}$ $min^{-1}$ |
| $ki_{f2}$ | Rate of BubR1 binding to KI2 motif | 1 $nM^{-1}$ $min^{-1}$ |
| $ki_{r1}$ | Rate of Bub1 dissociation from KI1 motif | 0.625 $min^{-1}$ |
| $ki_{r2}$ | Rate of BubR1 dissociation from KI2 motif | 0.625 $min^{-1}$ |
| $k_{scyc}$ | Rate of Cyclin B synthesis | 0.05 $nM{\cdot}min^{-1}$ |

| | | |
|---|---|---|
| $k_{dcyc}$ | Rate of Cyclin B degradation | 0.001 $min^{-1}$ |
| $k_{dcyc20}$ | Rate of APC:Cdc20 mediated degradation of Cyclin B | 0.004 $min^{-1}$ |
| $k_{ancyc}$ | Rate of Cdk1:CycB catalyzed activation of inactive eSAC | 1 $nM^{-1}$ $min^{-1}$ |
| $k_{incapp}$ | Rate of CAPP catalyzed inactivation of active eSAC | 1 $nM^{-1}$ $min^{-1}$ |
| $k_{imad20}$ | Rate of Cdc20 mediated inactivation of MCC complex | 0.01 $nM \cdot min^{-1}$ |
| $k_{dimcc}$ | Rate of dissociation of MCC complex | 1 $min^{-1}$ |
| $k_{acapp}$ | Rate of CAPP dephosphorylation | 0.1 $min^{-1}$ |
| $k_{icapp}$ | Rate of Cdk1 catalyzed phosphorylation of CAPP | 0.02 $nM^{-1} \cdot min^{-1}$ |
| $[Bub1]_T$ | Total concentration of Bub1 | 100 nM |
| $[BubR1]_T$ | Total concentration of BubR1 | 100 nM |
| $[CycB]_M$ | Concentration of Cyclin B in mitosis | 50 nM |
| $[Cdc20]_T$ | Total concentration of Cdc20 | 25 nM |
| $[Mad2]_T$ | Total concentration of Mad2 | 50 nM |
| $[CAPP]_T$ | Total concentration of CAPP (a Cdk1 counteracting protein phosphatase) | 50 nM |

**Table S3 Rates of MCC formation by eSAC according to its Bub1 and BubR1 binding state.** The four subscripts specify the Bub/BubR1 binding state of the four MELT repeats on an eSAC phosphodomain, using the rule: 0 → MELT, 1 → MELT:Bub1, 2 → MELT:Bub1:BubR1. For eSACs that bind 3 or more BubR1s (highlighted in red) cooperative interactions between SAC proteins is assumed and introduced through the multiplicative factor α = 2. The conversion rate by MELT:Bub1, $k_b$ = 0.1 $min^{-1}$ and by MELT:Bub1:BubR1, $k_{br}$ = 1 $min^{-1}$. The rates were multiplied by $8*10^{-3}$ to reproduce the dose-response curves shown in Figure 3E.

| Parameter | Value | Parameter | Value | Parameter | Value |
|---|---|---|---|---|---|
| $k_{0000}$ | 0 | $k_{1111}$ | $4*k_b$ | $k_{0201}$ | $k_b + k_{br}$ |
| $k_{1000}$ | $k_b$ | $k_{2000}$ | $k_{br}$ | $k_{0021}$ | $k_b + k_{br}$ |
| $k_{0100}$ | $k_b$ | $k_{0200}$ | $k_{br}$ | $k_{1120}$ | $2*k_b + k_{br}$ |
| $k_{0010}$ | $k_b$ | $k_{0020}$ | $k_{br}$ | $k_{1102}$ | $2*k_b + k_{br}$ |
| $k_{0001}$ | $k_b$ | $k_{0002}$ | $k_{br}$ | $k_{1210}$ | $2*k_b + k_{br}$ |
| $k_{1100}$ | $2*k_b$ | $k_{1200}$ | $k_b + k_{br}$ | $k_{1012}$ | $2*k_b + k_{br}$ |
| $k_{1010}$ | $2*k_b$ | $k_{1020}$ | $k_b + k_{br}$ | $k_{1201}$ | $2*k_b + k_{br}$ |
| $k_{1001}$ | $2*k_b$ | $k_{1002}$ | $k_b + k_{br}$ | $k_{1021}$ | $2*k_b + k_{br}$ |
| $k_{0110}$ | $2*k_b$ | $k_{0120}$ | $k_b + k_{br}$ | $k_{2110}$ | $2*k_b + k_{br}$ |
| $k_{0101}$ | $2*k_b$ | $k_{0102}$ | $k_b + k_{br}$ | $k_{0112}$ | $2*k_b + k_{br}$ |
| $k_{0011}$ | $2*k_b$ | $k_{0012}$ | $k_b + k_{br}$ | $k_{2101}$ | $2*k_b + k_{br}$ |

| | | | | | |
|---|---|---|---|---|---|
| $k_{1110}$ | $3*k_b$ | $k_{2100}$ | $k_b + k_{br}$ | $k_{0121}$ | $2*k_b + k_{br}$ |
| $k_{1101}$ | $3*k_b$ | $k_{2010}$ | $k_b + k_{br}$ | $k_{2011}$ | $2*k_b + k_{br}$ |
| $k_{1011}$ | $3*k_b$ | $k_{2001}$ | $k_b + k_{br}$ | $k_{0211}$ | $2*k_b + k_{br}$ |
| $k_{0111}$ | $3*k_b$ | $k_{0210}$ | $k_b + k_{br}$ | $k_{1112}$ | $3*k_b + k_{br}$ |
| $k_{1121}$ | $3*k_b + k_{br}$ | $k_{2102}$ | $k_b + 2*k_{br}$ | $k_{2211}$ | $2*k_b + 2*k_{br}$ |
| $k_{1211}$ | $3*k_b + k_{br}$ | $k_{2012}$ | $k_b + 2*k_{br}$ | $k_{2220}$ | $3*\alpha*k_{br}$ |
| $k_{2111}$ | $3*k_b + k_{br}$ | $k_{1220}$ | $k_b + 2*k_{br}$ | $k_{2202}$ | $3*\alpha *k_{br}$ |
| $k_{2200}$ | $2*k_{br}$ | $k_{0221}$ | $k_b + 2*k_{br}$ | $k_{2022}$ | $3*\alpha *k_{br}$ |
| $k_{2020}$ | $2*k_{br}$ | $k_{1202}$ | $k_b + 2*k_{br}$ | $k_{0222}$ | $3*\alpha *k_{br}$ |
| $k_{2002}$ | $2*k_{br}$ | $k_{0212}$ | $k_b + 2*k_{br}$ | $k_{1222}$ | $k_b + 3*\alpha *k_{br}$ |
| $k_{0220}$ | $2*k_{br}$ | $k_{1022}$ | $k_b + 2*k_{br}$ | $k_{2122}$ | $k_b + 3*\alpha *k_{br}$ |
| $k_{0202}$ | $2*k_{br}$ | $k_{0122}$ | $k_b + 2*k_{br}$ | $k_{2212}$ | $k_b + 3*\alpha *k_{br}$ |
| $k_{0022}$ | $2*k_{br}$ | $k_{1122}$ | $2*k_b + 2*k_{br}$ | $k_{2221}$ | $k_b + 3*\alpha *k_{br}$ |
| $k_{2210}$ | $k_b + 2*k_{br}$ | $k_{1212}$ | $2*k_b + 2*k_{br}$ | $k_{2222}$ | $4*\alpha *k_{br}$ |
| $k_{2201}$ | $k_b + 2*k_{br}$ | $k_{1221}$ | $2*k_b + 2*k_{br}$ | | |
| $k_{2120}$ | $k_b + 2*k_{br}$ | $k_{2112}$ | $2*k_b + 2*k_{br}$ | | |
| $k_{2021}$ | $k_b + 2*k_{br}$ | $k_{2121}$ | $2*k_b + 2*k_{br}$ | | |
